# The simulation extrapolation technique meets ecology and evolution: A general and intuitive method to account for measurement error

**DOI:** 10.1101/535054

**Authors:** Erica Ponzi, Lukas F. Keller, Stefanie Muff

**Affiliations:** Department of Evolutionary Biology and Environmental Studies, University of Zurich, Winterthurerstrasse 190, CH-8057 Zurich, Switzerland; Department of Biostatistics, Epidemiology, Biostatistics and Prevention Institute, University of Zurich, Hirschengraben 84, CH-8001 Zurich, Switzerland; Zoological Museum, University of Zurich, Karl Schmid-Strasse 4, CH-8006 Zurich, Switzerland

**Keywords:** Heritability, inbreeding coefficient, inbreeding depression, misassigned paternities, pedigree reconstruction, relatedness, SIMEX, uncertainty

## Abstract

1. Measurement error and other forms of uncertainty are commonplace in ecology and evolution and may bias estimates of parameters of interest. Although a variety of approaches to obtain unbiased estimators are available, these often require that errors are explicitly modeled and that a latent model for the unobserved error-free variable can be specified, which in practice is often difficult.
2. Here we propose to generalize a heuristic approach to correct for measurement error, denoted as simulation extrapolation (SIMEX), to situations where explicit error modeling fails. We illustrate the application of SIMEX using the example of estimates of quantitative genetic parameters, *e. g*. inbreeding depression and heritability, in the presence of pedigree errors. Following the original SIMEX idea, the error in the pedigree is progressively increased to determine how the estimated quantities are affected by error. The observed trend is then extrapolated back to a hypothetical error-free pedigree, yielding unbiased estimates of inbreeding depression and heritability. We term this application of the SIMEX idea to pedigrees “PSIMEX”. We tested the method with simulated pedigrees with different pedigree structures and initial error proportions, and with real field data from a free-living population of song sparrows.
3. The simulation study indicates that the accuracy and precision of the extrapolated error-free estimate for inbreeding depression and heritability are good. In the application to the song sparrow data, the error-corrected results could be validated against the actual values thanks to the availability of both an error-prone and an error-free pedigree, and the results indicate that the PSIMEX estimator is close to the actual value. For easy accessibility of the method, we provide the novel R-package `PSIMEX`.
4. By transferring the SIMEX philosophy to error in pedigrees, we have illustrated how this heuristic approach can be generalized to situations where explicit latent models for the unobserved variables or for the error of the variables of interest are difficult to formulate. Thanks to the simplicity of the idea, many other error problems in ecology and evolution might be amenable to SIMEX-like error correction methods.

## Introduction

Measurement error and other forms of uncertainty in variables of interest are commonplace in ecology and evolution, and there is thus a need for methods and practical tools to account for such errors in statistical models (see *e. g*. Solow, 1998; Macgregor et al., 2006; Reid et al., 2014; Steinsland et al., 2014; Wright et al., 2017; Mason et al., 2018). Measurement error can arise from countless sources in a wide range of studies, for example in the form of location error in telemetry and animal movement research (Montgomery et al., 2011; McClintock et al., 2014), error during the collection of phenotypic data (Hoffmann, 2000; Dohm, 2002; Macgregor et al., 2006; van der Sluis et al., 2010; Ge et al., 2017), misclassification in detection models and capture-recapture studies (Guillera-Arroita et al., 2014; Guelat and Kery, 2018), or error caused by spatial variability or uncertainty in the observation of climate variables (Bishop and Beier, 2013; Stoklosa et al., 2014) or biodiversity metrics (Haila et al., 2014; Mason et al., 2018).

When variables measured with error or estimated with uncertainty are used as explanatory variables in statistical analyses, parameter estimates may be biased (*e. g*. Fuller, 1987). To obtain unbiased parameter estimates, statistical models need to account for measurement error (see for example Gustafson, 2004; Carroll et al., 2006; Buonaccorsi, 2010, for an extensive treatment of frequentist and Bayesian measurement error correction techniques). Most standard measurement error correction methods require that the error model is known, that is, the error distribution and its parameters must be known from a quantitative assessment of the measurement error. In addition, some techniques require that latent (so-called “exposure”) models specify the distributions of the unobserved, true variables and possible dependencies on additional covariates, in particular when errors are modeled in a Bayesian framework (Muff et al., 2015; Ponzi et al., 2018). However, the error-generating mechanisms that blur true variables are often very complex and, hence, the distributions of unobserved, true variables are not always straightforward to specify. Consequently it can be difficult or even impossible to formulate and fit an explicit latent model that allows parametric measurement error modeling.

In cases where an explicit latent model for the unobserved, true variable is difficult to formulate, a general, heuristic method called simulation extrapolation (SIMEX) can be used to correct for measurement error. The SIMEX was originally introduced by Cook and Stefanski (1994) to correct for measurement error in continuous covariates of regression models, and later expanded to account for a broader range of regression models and error structures, such as misclassification error in discrete regression covariates and in the regression response (Kuechenhoff et al., 2006), or heteroschedastic error in covariates (Devanarayan and Stefanski, 2002). SIMEX is based on the rationale that more error leads to more bias in the estimated regression coefficients, and that progressively adding more error can reveal a pattern of the magnitude of the bias in dependence of the magnitude of the error. Based on this pattern, the algorithm extrapolates in the direction of less error, until the error-free estimate is reached. Thanks to its straightforward implementation without a latent model, its intuitive interpretation, and the possibility to cover a wide range of statistical models and error structures, SIMEX has been used extensively, with some applications also in ecology (*e. g*. Solow, 1998; Gould et al., 1999; Hwang and Huang, 2003; Melbourne and Chesson, 2006).

The main goal of this paper is to illustrate how the SIMEX approach can be further generalized to situations where it is not only difficult to specify the latent model, but also the error model, *i. e*. the distribution of the measurement error in the variables of interest. This occurs, for example, when the error mechanism acts not on the variable itself but on a lower level of the data. The example that we employ here, and which provided the initial motivation for this work, is error in estimates of inbreeding and relatedness(the variables of interest) resulting from errors in pedigrees (the data level where the error occurs). Pedigree error arises through various mechanisms, but in free-living organisms one of the major sources of pedigree error are incorrect paternities, when observed (social) behavior is used as a basis to assess parentage, but extra-pair paternities obscure the actual (genetic) relationships, leading to *misassigned paternities* (Keller et al., 2001; Griffith et al., 2002; Senneke et al., 2004; Jensen et al., 2007). These misassignments do not only affect the relatedness estimates of parents with their offspring, but all relatedness estimates among their descendants and their relatives. Thus, the correlation matrix of the similarity among relatives (the so-called relatedness matrix) is erroneous, as are the individual inbreeding coefficients. Consequently, we expect biased estimators for any quantitative genetic measure that builds on such information, such as estimates of heritability (Keller et al., 2001; Senneke et al., 2004; Charmantier and Reale, 2005), or of inbreeding depression (Keller et al., 2002; Visscher et al., 2002; Reid et al., 2014).

While it is difficult to formulate an explicit parametric error model for the variables of interest, *i. e*. inbreeding and relatedness coefficients and the relatedness matrix in the presence of misassigned paternity error in the pedigree, it is relatively straightforward to simulate error at the pedigree level and to repeatedly estimate the quantitative genetic measures with different levels of error. This is where the SIMEX idea enters: Instead of increasing the error variance of a continuous covariate as in the traditional SIMEX, we successively increase the proportion of misassigned paternities in the pedigree to obtain information about the bias in quantitative genetic estimates as the pedigree error increases. In a second step, the observed trend upon increasing the error proportion is extrapolated back to that of an error-free pedigree. This novel method, which we denote pedigree-SIMEX (PSIMEX), is straightforward to apply, given that the the proportion of misassigned paternities, as well as their distribution in the actual pedigree (*e. g*. proportions varying over time) are known.

Here we test the validity of the PSIMEX approach with different simulated pedigree topologies, and show that the method can substantially reduce or eliminate the bias in estimates of heritability and inbreeding depression. We then apply the PSIMEX algorithm to an empirical data set from a population of song sparrows, where apparent paternities (observed from social behavior) and actual (genetic) paternities are not always corresponding. Since paternities were determined both socially and genetically in this population, we were able to compare the PSIMEX estimates of heritability and inbreeding depression derived from the apparent pedigree to the estimates derived from the actual pedigree. Our application to the song sparrow data suggests that the PSIMEX method performs well not only in simulations but also with real field data. To facilitate the use of PSIMEX, we provide the novel R-package `PSIMEX` (Ponzi, 2017).

## Theory

### The original SIMEX algorithm

We start by outlining how SIMEX works in its simplest form, as originally proposed by Cook and Stefanski (1994). Assume that a continuous variable of interest *x* is blurred by *classical additive measurement error*, such that only *w* = *x* + *u* can be observed, where the measurement error *u* is assumed to be independent of *x* and distributed as 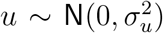 with error variance 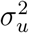. Further assume that *w* instead of the unobservable *x* is used as a covariate in a simple linear regression model, *y* = *α* + *β_w_w* + *ϵ*. This is a typical measurement error, or errors-in-variables problem, known to lead to a biased regression parameter estimate, whenever 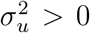 (Fuller, 1987; Carroll et al., 2006). Using 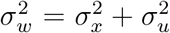 and the assumption that the error *u* is independent of *x*, it is quite straightforward to see that the error-prone regression parameter *β_w_* is an estimator of

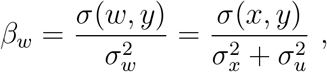

which is less than the true slope *β_x_* by an *attenuation* factor 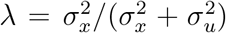, so that *β_w_* = λ*β_x_*. Although this suggests that measurement error will generally lead to underestimated effect sizes, this is only true in this very simple situation. Even relatively standard regression models may yield upwardly biased parameter estimates, for example when an error-prone covariate is correlated with another covariate, in the presence of interactions, or in logistic regression (Carroll et al., 2006; Freckleton, 2011; Muff and Keller, 2015).

To obtain estimates of the true slope *β_x_* instead of the biased *β_w_*, the SIMEX algorithm is based on the heuristic that more error will generally lead to more bias. By systematically increasing the error in a simulation (SIM) step and then extrapolating (EX) the pattern of change in parameter estimates with increasing error backward, one approximates the parameter that one would obtain if there was no error in the data. Figure 1 depicts the SIMEX idea graphically. In the case introduced above of classical additive measurement error in a continuous covariate, the error variance 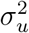 is artificially increased by adding more random error to the covariate of interest. For each error level (*i. e*. each predefined increase of the error variance), the procedure is iterated *K* times and regression parameters and standard errors are estimated and stored for each iteration. In the extrapolation phase, the observed trend upon increasing error is extrapolated in the direction of less error, and an error-corrected SIMEX estimate is obtained for zero error. The extrapolation function can have different forms, depending on the trend which is observed as the error increases. The most common extrapolation functions are linear and quadratic, but polynomial or nonlinear functions are also sometimes used. Standard errors for the error-corrected estimate are obtained by summing over two sources of variability, the sampling error of the *K* simulations and the standard errors that are obtained from each regression. Details about the computation of the standard error in the SIMEX algorithm are given in section 1 of Appendix 1. In addition, the reader is referred to Stefanski and Cook (1995) and Apanasovich et al. (2009).

**Figure 1:**
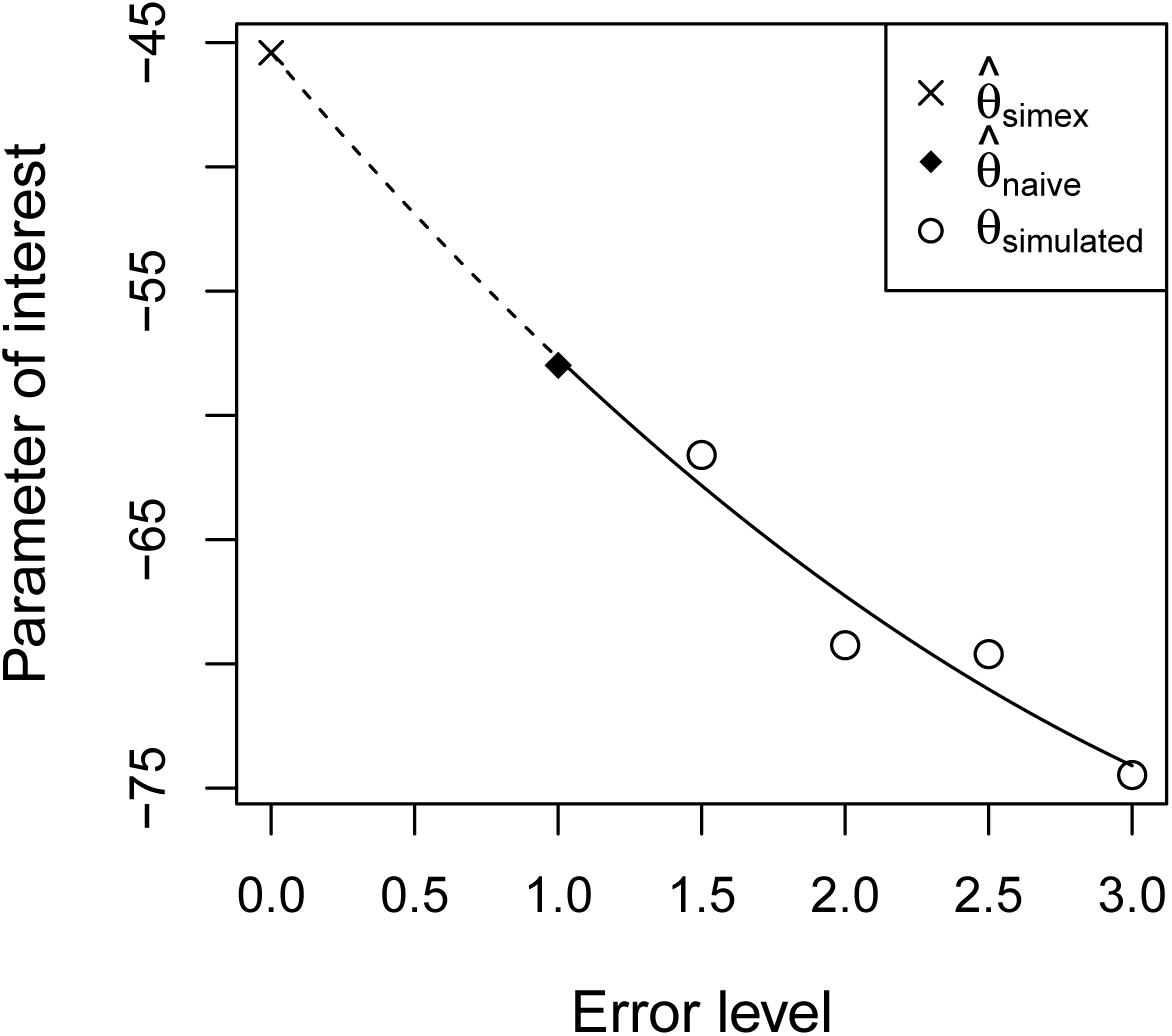
Illustration of the SIMEX procedure. The error level is increased in predefined interval steps, the parameter of interest is estimated at each error level, and a function is fitted on the observed trend upon increasing error. An error-corrected estimate is obtained by extrapolating the function back to an error level of zero.

To apply the SIMEX algorithm and to know when zero error is reached, the initial error *model* (*e. g*. 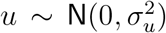) and error model *parameter* (*e. g*. the value of 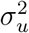) must be known. The error model defines the mechanism according to which more error must be generated in the simulation phase, and the parameter value defines the fixed point from where back-extrapolation begins. Incorrect assumptions about the error model or error parameter may result in a biased simulation mechanism and an incorrect extrapolation function and, therefore, yield parameter estimates that are over or under-corrected. In the worst case, correction may be in the wrong direction. Thus, to use SIMEX – or, indeed, any measurement error correction technique – it is crucial to have a good understanding of the error model and good estimates of the error parameters through repeated measurements or validation data. In the majority of applications, the SIMEX method results in a substantial bias reduction in the estimators, although there are some particular cases where the bias reduction is not considered sufficient and other methods might be preferable (*e. g*. Fung and Krewski, 1999; Hwang and Huang, 2003; Carroll et al., 2006, pp. 121 – 123).

### Extensions of SIMEX

Since the original contribution by Cook and Stefanski (1994), SIMEX has been extended to account for different types of errors and models. Examples include non-additive error models (Eckert et al., 1997) or heteroschedastic error variances (Devanarayan and Stefanski, 2002). Another extension is the so-called misclassification SIMEX (MCSIMEX), which accounts for error in discrete variables by simulating data with higher misclassification probabilities and extrapolating back to no misclassification. These extensions highlight the flexibility of SIMEX in situations where standard error modelling procedures are challenging to implement. The extension we suggest here covers the case when the error mechanism does not act on the variable itself, but on an underlying structure, illustrated on error in the pedigree of a study population.

### SIMEX for pedigrees: PSIMEX

We describe the SIMEX idea applied to pedigree error, denoted as PSIMEX. The procedure starts from an initial error level *ζ_I_* ∈ (0,1), *i. e*. the proportion of misassigned parentages that occur in the pedigree, for example *ζ_I_* = 0.1, and randomly generates additional misassigned parents, resulting in error levels *ζ* > *ζ_I_*, for example *ζ* = 0.2, 0.3, 0.4 and so on. To achieve a desired error proportion *ζ* > *ζ_I_* of misassigned parents, we cannot simply pick a proportion of *ζ* – *ζ_I_* parents and replace them, because this may include parents that were already incorrectly assigned, so that the effective proportion of error would be too low. To account for this circumstance, the actual proportion of additional misassignments needed at each step is calculated from the equation

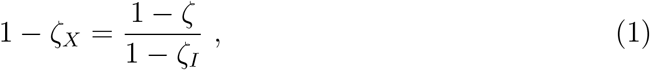

where *ζ_X_* is the *effective* error proportion that has to be added to obtain a nominal error level of *ζ*. As an example, assume that the initial parentage error rate is *ζ_I_* = 0.17 and that the aim is to increase this proportion to *ζ* = 0.30 (0.17+0.13). If only a proportion of 0.13 of the parental relations is randomly picked and reassigned, it is expected that some parents that were already misassigned are randomly misassigned again. The effective error proportion is then less than 0.30. In fact, equation (1) shows that we need to pick and re-assign 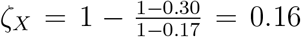 of the parents to obtain an expected error proportion of 0.30.

Each randomly selected parent is then replaced by connecting the offspring to another individual according to a known error-generating mechanism (*e. g*. some parents might be more likely to be chosen than others, see below). For each error proportion *ζ* the procedure is repeated a fixed number of *K* (*e. g*. 100) times, and for each *k* = 1, 2,…, *K*, the pedigree is recalculated including the newly misassigned connections and estimated quantities of interest, denoted as 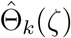, are stored. The algorithm then averages over the *K* values of 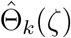 to obtain the desired estimate at error proportion 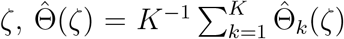. Given that these means are calculated for a sequence of values *ζ* > *ζ_I_*, it is possible to estimate a functional dependence between 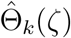 and *ζ*. The estimate at zero error 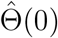, obtained by extrapolating in the direction of decreasing error, corresponds to the error-corrected estimate denoted as 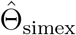. Different extrapolation functions, as well as criteria to select it, are reported in the following sections of the manuscript.

When applying PSIMEX, precise knowledge of the initial error proportion *ζ_I_* and the mechanisms leading to this value are necessary. An intrinsic assumption of the simulation phase, when additional error is generated, is that the same error-generating mechanism is used as in the observed data. As an example, the percentage of misassigned parents may fluctuate over time with fluctuating population size or sex ratio. In many situations, misassignments might affect solely fathers and occur within the same generation, so it might be logical to assume replacement of only fathers with random individuals chosen from the same generation. Such structural aspects of the error-generating mechanisms must be taken into account in the PSIMEX procedure, because the relationship between error and bias may otherwise not reflect the true trend. Examples of the effects of incorrect assumptions about the initial error proportion or the error structure are given in sections 4 and 5 of Appendix 1.

### Quantitative genetic measures

To illustrate how the PSIMEX idea works, we apply it to the estimation of two important quantitative genetic parameters, heritability and inbreeding depression. Heritability quantifies the proportion of phenotypic variance in a trait that is due to additive genetic factors and is key to predicting the response to selection (Lynch and Walsh, 1998). Inbreeding depression quantifies the reduction in fitness of offspring resulting from matings among relatives and is key to understanding mating system evolution and dispersal (Keller and Waller, 2002; Charlesworth and Willis, 2009).

Both quantities can be estimated by fitting (generalized) linear mixed models to phenotypic or fitness data, using the so called *animal model* (Henderson, 1976; Lynch and Walsh, 1998; Kruuk, 2004). For a continuous trait with multiple measurements per individual the animal model can be written as

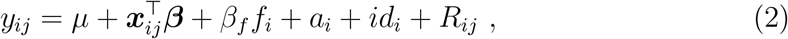

where *μ* is the population mean, ***β*** is a vector of fixed effects, ***x***_*ij*_ is the vector of covariates for individual *i* at the *j*^th^ measurement occasion, *f_i_* is the inbreeding coefficient of individual *i*, which reflects how related an animal’s parents are, and *β_f_* is the fixed effect of inbreeding depression (Lynch and Walsh, 1998). The last three components of equation (2) are random effects, namely the additive genetic effects (or breeding values) *a_i_* with dependency structure 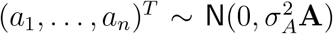, the independent effects for the animal identity 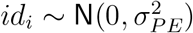 accounting for the permanent environmentally-induced differences among individuals, and an independent Gaussian residual term 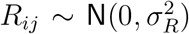 that captures the remaining (unexplained) variability. The dependency structure of the breeding values *a_i_* is given by the additive genetic relatedness matrix **A** (Lynch and Walsh, 1998), which is traditionally derived from the pedigree. In this framework, the narrow-sense heritability *h*^2^ is defined as

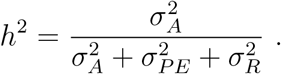

Inbreeding depression is estimated by the regression coefficient *β_f_*, with a negative slope indicating inbreeding depression.

Not all components of the animal model (2) are always necessary. For example, the breeding values *a_i_* are often omitted when estimating inbreeding depression, although this can result in biased estimates of inbreeding depression and is thus not recommended (Reid et al., 2008; Becker et al., 2016). Also, inbreeding *f_i_* is not always included as a covariate when estimating heritability, although it is generally recommended to account for potentially higher phenotypic similarity between individuals with similar levels of inbreeding (Reid et al., 2006; Reid and Keller, 2010), but also because ignoring the effects of inbreeding may lead to biases in estimates of the additive genetic variance 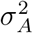 (Wolak and Keller, 2014). On the other hand, model (2) can be further expanded to include additional variance components, such as the maternal variance 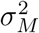, or nest or time effects (Kruuk and Hadfield, 2007; Wilson et al., 2010). Moreover, it can be formulated as a generalized linear mixed model (GLMM), for example for binary traits. We will use a GLMM in our application to the song sparrow data below.

In the presence of misassigned parentages in the pedigree structure, the estimates in the relatedness matrix **A** will necessarily suffer from error, which, in turn, may bias the estimates of the variance components and thus the estimates of heritability *h*^2^. Similarly, the accurate quantification of inbreeding coefficients *f_i_* will be hampered by misassigned parentages in the pedigree, which will result in biased estimates of inbreeding depression *β_f_*. Parametric assumptions about the error mechanism that distorts relatedness and inbreeding estimates can be challenging to formulate, because the error propagates from the pedigree structure. The resulting error distribution is therefore expected to be non-standard, as illustrated in Appendix 1, Section 2.1. This is exactly where the PSIMEX algorithm is useful, because it is possible to simulate error in *f_i_* and in **A** by simulating error in the pedigree, where it is more straightforward to specify an error model.

## Simulation study

To illustrate and test the PSIMEX procedure we carried out a simulation study with a set of simulated pedigrees, generated using the function `generatePedigree()` from the R package `GeneticsPed` (Gorjanic and Henderson, 2007). We then introduced increasing proportions of misassignments of fathers, which we replaced with a randomly chosen male individual from the same generation, using a constant misassignment probability across all generations in the pedigree. After quantifying the effects of the errors on estimates of heritability and inbreeding depression, we applied the PSIMEX procedure to obtain error-corrected estimates of the two measures, starting from different initial error proportions.

### Effects on heritability

To understand the effect of pedigree error on heritability, we simulated phenotypic traits *y_i_* as

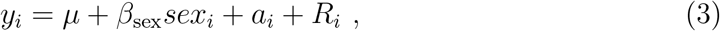

with *μ* = 10, *β*_sex_ = 2, 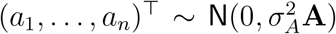 and independent 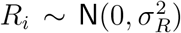, using 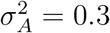 and 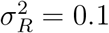. Breeding values *a_i_* were generated with the `rbv()` function from the R package `MCMCglmm` (Hadfield, 2010) that accounts for the dependency structure given by the relatedness matrix **A** from the pedigree. The simulated heritability was thus 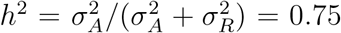, and estimates were obtained by fitting model (3) using the `MCMCglmm` package and extracting posterior means and variances of the `MCMC` samples. Note that the same animal model could be implemented using *e. g*. integrated nested Laplace approximations (INLA, Rue et al., 2009; Holand et al., 2013), although we used `MCMCglmm` throughout this paper.

A total of 50 different pedigrees were generated, with 50 mothers and 50 fathers in each of 30 generations. For each pedigree, we simulated error proportions *ζ* of paternal misassignments ranging from 0 to 1.0 in steps of 0.1, and the procedure was repeated 100 times for each error level and each pedigree. At each iteration, the naive *ĥ*^2^ was calculated together with its standard error. Following Wilson et al. (2010, Supplementary File 5), inverse gamma priors were used for all variance components, namely 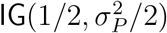, where 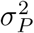 is the total phenotypic variance of the trait, but the results were not sensitive to the prior choice. For example, we obtained the same results when assigning 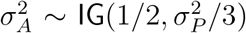 and 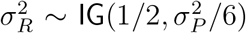, which reflects a prior belief that a larger proportion of variance is captured by the additive genetic component rather than by the environment.

### Effects on inbreeding depression

Given that the relatedness structure and inbreeding in a population depend – among other things – on the effective population size and the variance in reproductive success (Reid and Keller, 2010), it seemed likely that the structure and topology of the pedigree influence the effects of pedigree error on inbreeding depression. We therefore considered different pedigree structures that induce different levels of inbreeding in a population. Different pedigree structures were simulated through different *Ne/Nc* ratios, defined as the ratio of the effective population size (*Ne*) to the census size (*Nc*) (Frankham, 1995; Palstra and Fraser, 2012). Different *Ne/Nc* ratios were obtained by changing the number of reproductive males and females per generation, while keeping the population size constant at 100. In the first case, both numbers of fathers and mothers per generation were set to 50, which led to *Ne/Nc* = 1, so that all individuals had offsprings. As a consequence, the average degree of relatedness among individuals was comparatively low, implying a low average inbreeding coefficient throughout generations. The second case corresponded to pedigrees with only 15 mothers and fathers per generation (*Ne/Nc* = 0.3) and the third case to pedigrees with only 5 mothers and fathers per generation (*Ne/Nc* = 0.1). As the 100 individuals of a generation thus originated from only 30 or 10 parents, respectively, the mean relatedness among individuals increased, and their matings thus led to more highly inbred offspring. Note that this change in pedigree structure does not affect heritability estimates, which only depend on the initial variance components and not on the topology of the pedigree.

Individual inbreeding coefficients (*f_i_*) were derived from the pedigree using the `calcInbreeding()` function from the R package `pedigree` (Coster, 2013). Fitness traits *y_i_* for individual *i* were then simulated according to

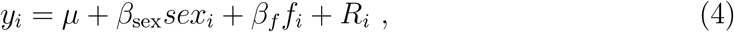

with a population mean of *μ* =10, sex effect of *β*_sex_ = 2, inbreeding depression *β_f_* = −7, and a residual term *R_i_* ~ `N`(0, 0.1). For each of the three pedigree topologies, we generated 100 distinct error-free pedigrees. Error was then added, ranging from *ζ* 0 to 1.0 in steps of 0.1. A total number of 100 pedigrees were simulated for each error level and each originally generated pedigree. In each iteration, the linear regression model (4) was fitted, but using the inbreeding coefficients *f_i_* that were estimated from the pedigree with misassigned paternities as covariates. The estimated inbreeding depression *β_f_* was stored in each iteration.

### Application of PSIMEX to simulated pedigrees

To assess the performance of the PSIMEX algorithm in recovering error-free estimates of heritability and inbreeding depression, we started with initial error proportions of *ζ_I_* = 0.1, 0.2, 0.3 and 0.4 and applied PSIMEX to all the simulated original pedigrees. For each error proportion *ζ_I_*, we selected one erroneous pedigree out of the 100 simulated ones and provided the algorithm with the known initial error proportion and correct assumptions about the error mechanism, *i. e*. obtained by random replacement of fathers with male individuals from the same generations and with constant mis-assignment probability across all generations in the pedigree. During the simulation phase where error was increased, estimates were averaged across *K* = 100 iterations of the same error level and stored together with their standard errors. Error-corrected estimates 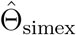 for *h*^2^ and *β_f_* were obtained by fitting a linear, a quadratic and a cubic extrapolation function to the trend upon increasing error proportions. The function with lowest AIC was retained. The results were compared to the corresponding naive estimates 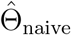, which were obtained using the error-prone estimates of *f_i_* from each original pedigree. The code for all simulations is given in Appendices 2 and 3.

### Simulation results

#### The effects of pedigree error on heritability

Misassigned paternity error caused a clear decreasing trend in heritability, with a continuous decline in the estimate as the error proportions increased (Figure 2). This is a consequence of the decrease in the estimates of 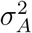 due to the increasing pedigree error, which causes information loss in the relatedness matrix **A**.

**Figure 2:**
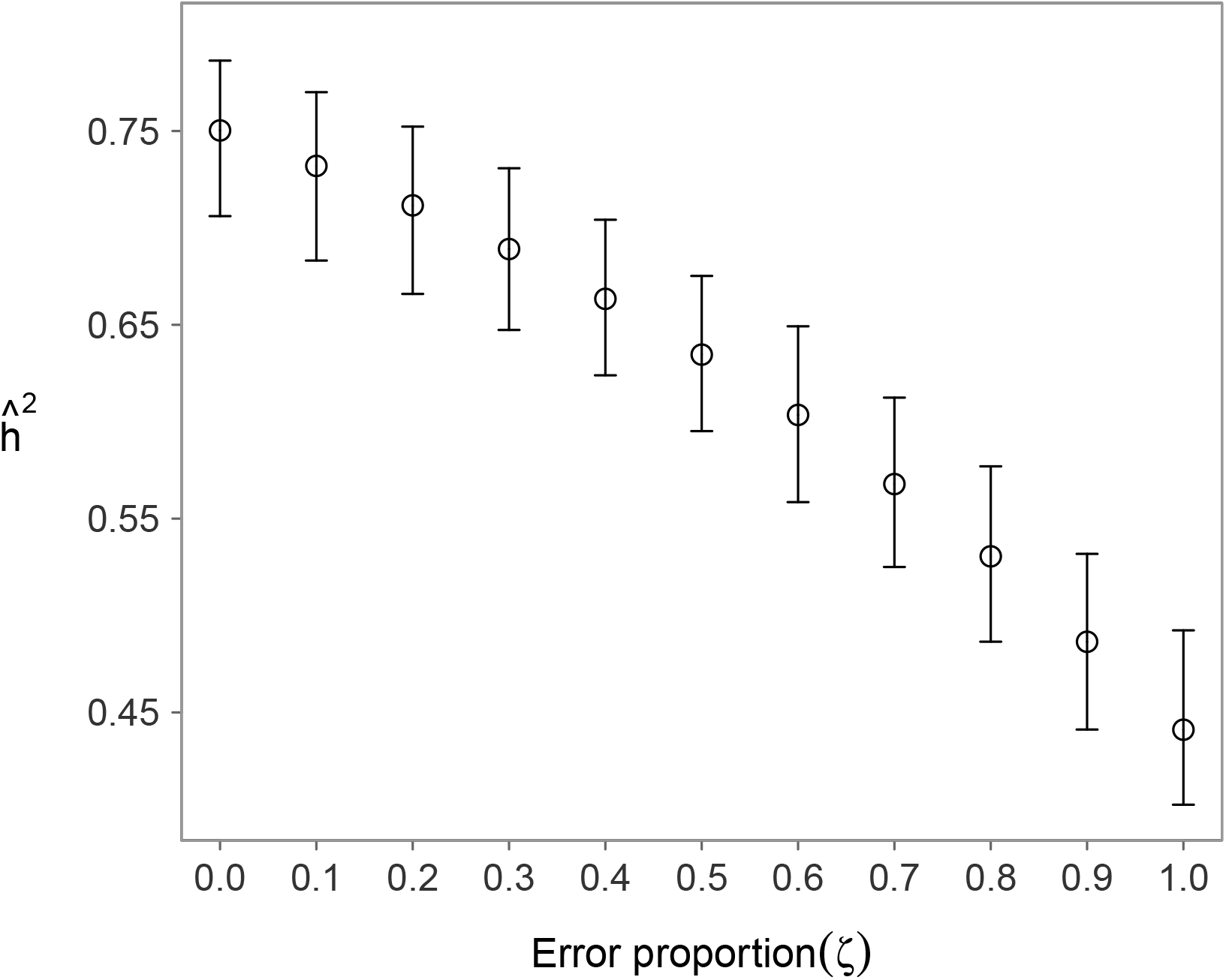
Effect of increasing the paternity error proportion on heritability in simulated pedigrees. Mean estimates of heritability *ĥ*^2^ from 100 simulations for each pedigree with increasing error rates *ζ* = 0.1,…, 1 are shown with their 5% to 95% sample quantile intervals at each error proportion. A clear decreasing trend is observed in the estimate as the error proportion increases.

In our simulations, all mothers were always correctly assigned to their offspring. Thus, even when all paternities were assigned randomly (*ζ* = 1), we do not expect heritability estimates of *h*^2^ = 0, since at least half of the parent-offspring pairs in the pedigree were still correct. With all mothers assigned correctly and all fathers assigned incorrectly, one would expect heritability estimates to equal half the true heritability of 0.75, *i. e. h*^2^ = 0.375 (the heritability estimate from a mother-offspring regression, Lynch and Walsh (1998)). However, for complete paternal misassignments (*ζ* = 1) we obtained an average heritability of *h*^2^ = 0.44 (95% quantile interval from 0.40 to 0.49). This is higher than expected, because the misassigned fathers were, on average, still related to the true fathers due to the small population size (100 individuals per generation). We tested this explanation for the higher than expected heritability estimates with *ζ* = 1 by increasing the number of individuals in each generation to 1000 in 50 pedigrees. This reduced the relatedness between true and randomly assigned fathers and resulted in an average heritability estimate close to expectations (*h*^2^ = 0.37, 95% quantile interval from 0.34 to 0.40).

#### The effects of pedigree error on inbreeding depression

As expected, the trend in estimates of inbreeding depression (*β_f_*) depended on the structure of the simulated pedigree (Figure 3). Adding misassigned paternities to pedigrees with *Ne/Nc* = 1 (low average inbreeding) led to underestimation of in-breeding depression, with a trend towards zero as error proportions *ζ* increased (Figure 3a). An opposite pattern was observed in pedigrees with *Ne/Nc* = 0.1 (high average inbreeding), where the estimated inbreeding depression became stronger with increasing paternity error (Figure 3c). Intermediate pedigrees with *Ne/Nc* = 0.3 showed an initial trend of increased inbreeding depression, followed by a decrease and a trend towards zero, resulting in a U-shaped pattern (Figure 3b). The difference of these patterns can be due to the fact that randomly replacing fathers in a pedigree structure that is characterized by high initial inbreeding (*i. e. Ne/Nc* = 0.1) causes, on average, a decrease in the inbreeding value, while when the initial inbreeding value is on average low (*i. e. Ne/Nc* = 1), a random replacement of fathers leads to generally more inbred breeding pairs, that is, a higher level of inbreeding in the population. Consequently, pedigree errors may affect the absolute value of inbreeding depression in opposite directions.

**Figure 3:**
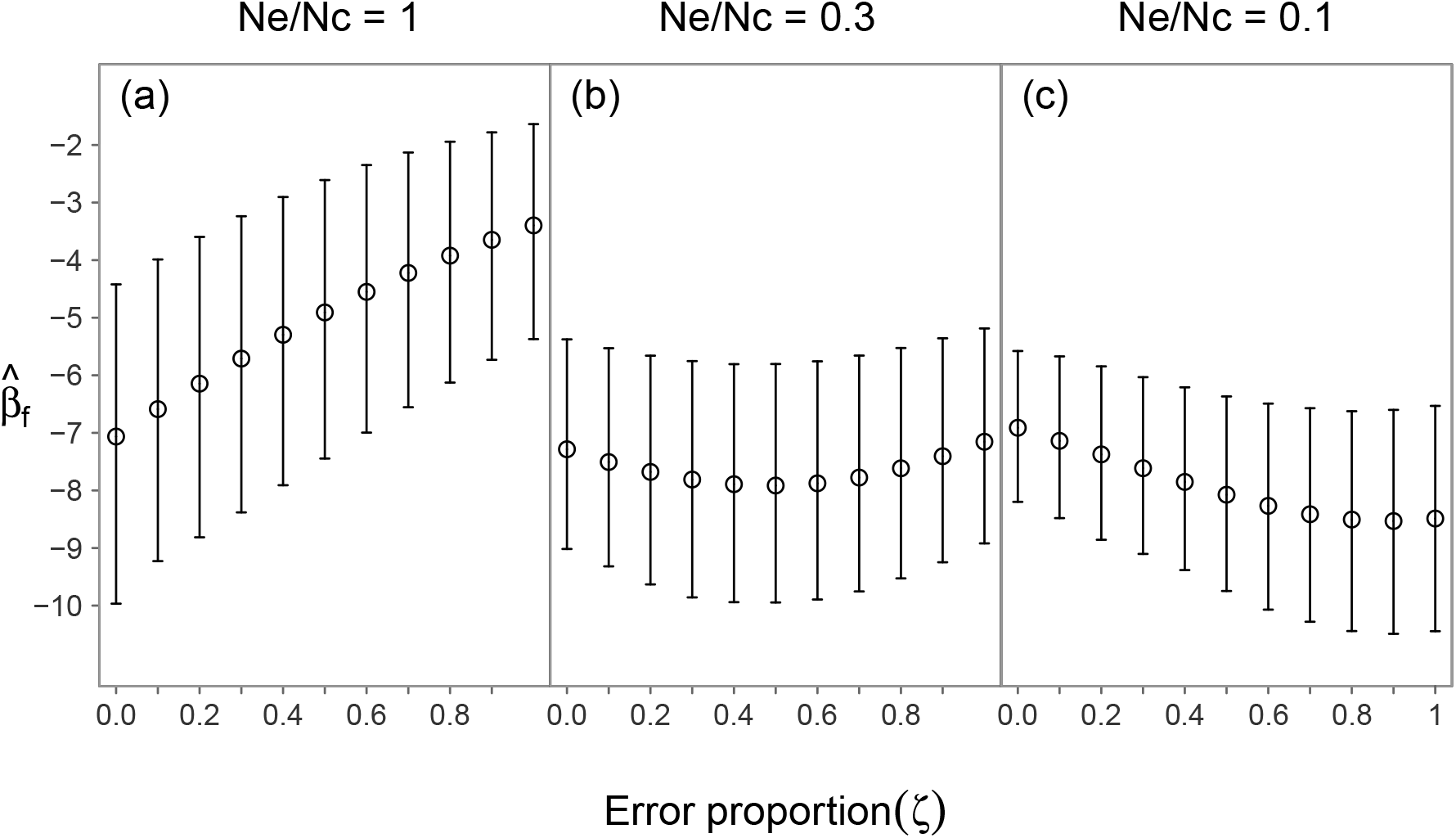
Effects of increasing paternity error proportion on inbreeding depression in simulated pedigrees with each of the three different pedigree topologies (*N_e_/N_c_* = 1, 0.3, and 0.1). Mean estimates of inbreeding depression 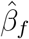 from 100 simulations with increasing error rates *ζ* = 0.1,…, 1 are shown with their 5% to 95% sample quantile intervals at each error proportion.

#### PSIMEX estimates of heritability and inbreeding depression

The PSIMEX procedure yielded heritability estimates that were much closer to the simulated true value of *h*^2^ than the naive estimates obtained ignoring the error (Figure 4, left). PSIMEX recovered essentially consistent estimators, with increasing uncertainty for larger values of *ζ_I_*. Even when starting from an initial error proportion as high as 0.4, the 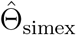 estimates were still much closer to the actual value than 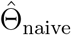, although with larger uncertainty than for smaller initial values. In most cases (56%) the quadratic extrapolation function had the smallest AIC and was thus chosen to obtain the PSIMEX estimate. In the remaining cases, the cubic function was selected.

**Figure 4:**
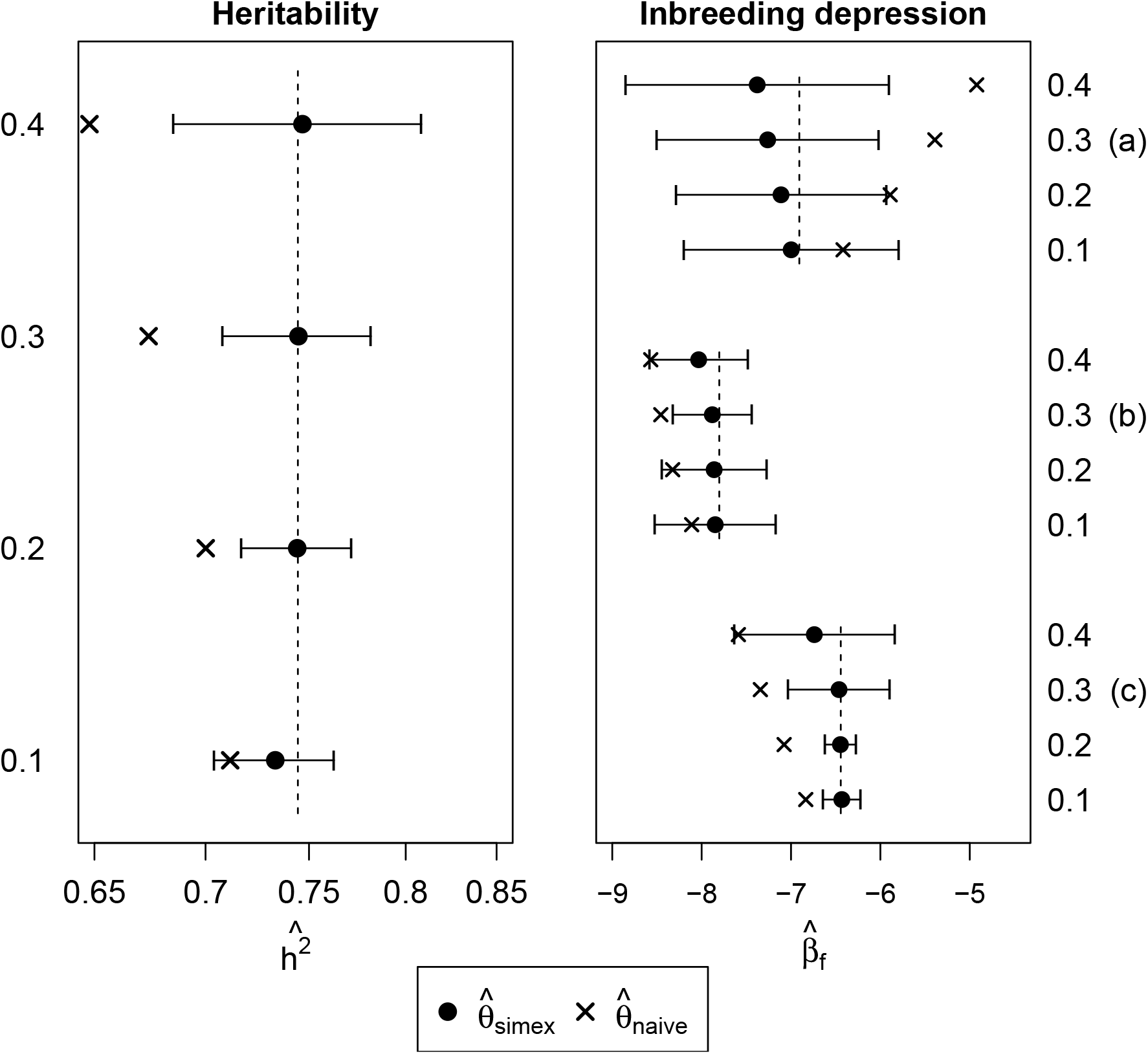
Naive 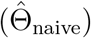 and error-corrected 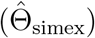 estimates of heritability and inbreeding depression from the simulated pedigrees. For the error-corrected PSIMEX estimates, means and 95% confidence intervals from the simulations for PSIMEX estimates are reported, whereas only the point estimates are shown for the naive results. The dashed line represents the actual simulated value for the respective pedigree. Four different estimates are given, corresponding to four initial error proportions (*ζ_I_* = 0.1, 0.2, 0.3 and 0.4). For inbreeding depression, results are reported for pedigree topologies a) *Ne/Nc* = 1, b) *Ne/Nc* = 0.3 and c) *Ne/Nc* = 0.1.

PSIMEX also yielded estimates of inbreeding depression 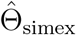 that were much closer to the simulated true value for all three pedigree topologies (*Ne/Nc* = 1.0, 0.3, 0.1) and all four initial error proportions (*ζ_I_* = 0.1, 0.2, 0.3, 0.4; Figure 4). Even though 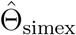 estimates were somewhat less consistent for larger *ζ_I_*, because it is then more difficult to recover an appropriate extrapolation function, they were still much closer to the true simulated values than the corresponding naive estimates obtained ignoring the error. In 100%, 50% and 79% of the cases for pedigrees with *Ne/Nc* = 1, *Ne/Nc* = 0.3, and *Ne/Nc* = 0.1, respectively, the cubic extrapolation function had the smallest AIC and was thus chosen, whereas a quadratic extrapolation function was selected in the remaining cases.

## Empirical example: Song sparrows

### Study population

We also applied the PSIMEX approach to an empirical study of a population of free-living song sparrows (*Melospiza melodia*) on Mandarte Island, Canada. Over more than 40 years, 31 generations of song sparrows have been monitored, yielding a pedigree of 6095 individuals, together with data on morphological and life history traits. For details on data collection and methods see Smith and Keller (2006).

Thanks to the small population size (roughly 30 breeding pairs), pedigree data are essentially complete. However, extra-pair matings are common (*e. g*. Reid et al., 2014), leading to pedigree error in the so-called *apparent pedigree* when fathers are inferred from observing parents providing parental care. The apparent pedigree contains misassigned paternities, but not misassigned maternities, since all mothers are the genetic parent of the offspring they feed (Reid et al., 2014; Germain et al., 2016). Genetic information from 13 microsatellite loci has been used to assign each offspring its genetic father, yielding an almost exact reconstruction of the *actual pedigree* (Sardell et al., 2010). A comparison of the apparent and actual pedigree yields an estimate of the percentage of misassigned paternities of 0.17. Note that this percentage differs from extra-pair paternity rates reported elsewhere, because we restricted the analysis to individuals with records of tarsus length and juvenile survival.

The circumstance that we have both an error-prone and an essentially error-free version of the same pedigree renders the song sparrows an ideal study system to test the PSIMEX algorithm.

### PSIMEX for the song sparrow analyses

Our analyses focused on the heritability of tarsus length, which has previously been estimated to be approximately *h*^2^ = 0.32 among Mandarte song sparrows (Smith and Zach, 1979), and on juvenile survival, which is known to exhibit inbreeding depression in this population (*e. g*. Reid et al., 2014). We applied PSIMEX to obtain error-corrected estimates of the heritability of tarsus length and the magnitude of inbreeding depression in juvenile survival from the apparent (error-prone) pedigree with a paternity error rate of *ζ_I_* = 0.17. We assumed this proportion to be constant across generations, and that misassigned fathers were randomly selected from the same generation. We assumed that matings were random, despite some weak indication that relatively inbred animals might be slightly more likely to mate with closely related individuals (Reid et al., 2006). The corresponding estimates from the actual (error-free) pedigree served as a benchmark for the error-corrected estimates 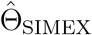. To estimate heritability of tarsus length we fitted the model

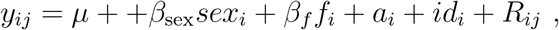

where *y_ij_* is the *j*^th^ measurement of tarsus length for individual *i*, and *sex_i_* and inbreeding coefficient *f_i_* are fixed effects. Random effects were as defined in equation (2). The model was fitted with `MCMCglmm` and posterior means and variances were extracted from the `MCMC` samples. Inverse gamma priors 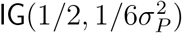 were used for the three variance components 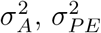 and 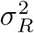. On the other hand, juvenile survival *y_i_* is a binary trait indicating survival of individual *i* from an age of six days to one year (1=yes, 0=no), thus we used a GLMM. The binary survival variable *y_i_* can be interpreted as the realization of a slow count process, thus a complementary log-log (cloglog) link was used, which encodes for `P`(*k* = 0) of a Poisson model (McCullagh and Nelder, 1989, pp. 151-153). The model was thus given as

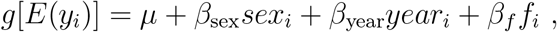

where *E*(*y_i_*) = *p_i_* is the Bernoulli distributed survival probability of individual *i*, and *g* is a complementary log-log link function. Besides the sex of the individual, we also included the year a bird was born as a categorical covariate. More complex models can be formulated, for example including an interaction term between sex and inbreeding coefficient (see Reid et al., 2014), but we ignored these extensions for the purpose of this paper. The model was fitted in a likelihood framework using `lme4` in `R`, as in Reid et al. (2014).

In both the heritability and the inbreeding depression analysis, we set the number of iterations in the PSIMEX procedure to *K* = 100. Linear, quadratic and cubic functions were used to describe the trend of the estimate of interest and to extrapolate a value for the PSIMEX estimate 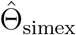. The code for the application of PSIMEX to the song sparrow data is given in Appendices 6 and 7.

### Song sparrows results

The error-corrected PSIMEX estimates given by the best fitting (*i. e*. minimum AIC) extrapolating functions were closer to the actual values obtained with the genetically correct actual pedigree than the corresponding naive estimates, both for heritability and for inbreeding depression (Table 1). Figure 5 shows both the trend of the simulated values for increasing error proportions and the extrapolated values for zero error given by PSIMEX. Both for heritability and inbreeding depression, the AIC-criterion suggested that a quadratic extrapolation function fitted the error trend best.

**Figure 5:**
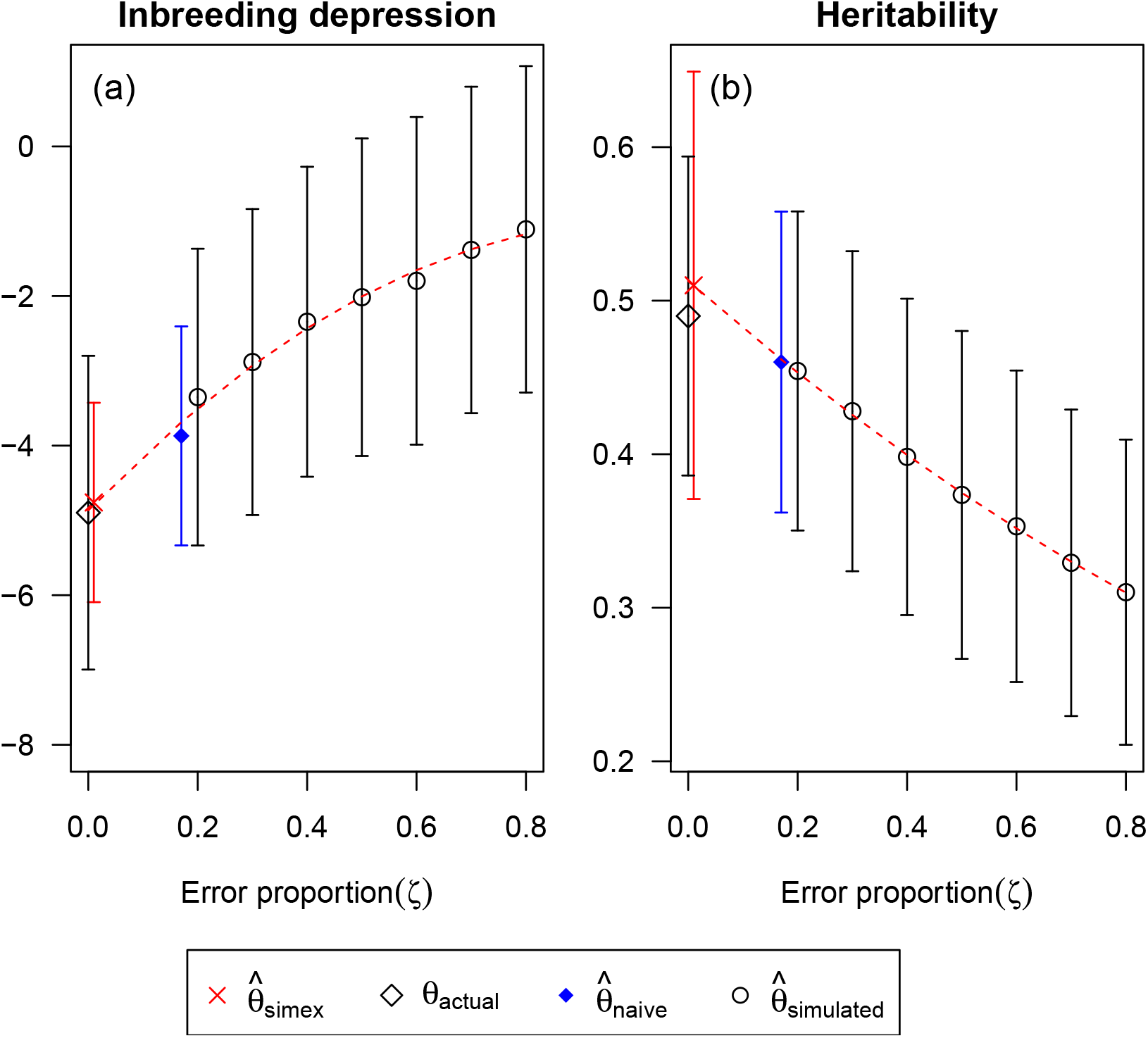
Results of the PSIMEX procedure when error-correcting heritability of tarsus length (a) and inbreeding depression of juvenile survival (b) in the song sparrow dataset. The initial error proportion was *ζ* = 0. 17. The trend upon increasing error proportions is shown together with the extrapolated values obtained from the best extrapolation function (quadratic). The naive and the actual estimates from the genetic pedigree are also shown. PSIMEX estimates are much closer to the actual values than the naive estimates both for heritability and inbreeding depression.

**Table 1:** Estimates of heritability (*ĥ*^2^) of tarsus length and inbreeding depression 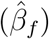 of juvenile survival in the song sparrow dataset. Actual, naive and PSIMEX estimates are reported together with their 95% confidence intervals (CIs). The quadratic function was chosen as the best extrapolating function (lowest AIC).

| | $\hat{h}^2$ | 95% CI | $\hat{\beta}_f$ | 95% CI |
| --- | --- | --- | --- | --- |
| $\hat{\theta}_{\text{actual}}$ | 0.49 | 0.38 to 0.60 | −4.90 | −6.26 to −3.53 |
| $\hat{\theta}_{\text{naive}}$ | 0.46 | 0.36 to 0.56 | −3.87 | −5.33 to −2.40 |
| $\hat{\theta}_{\text{simex}}^{(\text{linear})}$ | 0.50 | 0.39 to 0.61 | −4.38 | −6.63 to −1.92 |
| $\hat{\theta}_{\text{simex}}^{(\text{quadratic})}$ | **0.51** | **0.38 to 0.66** | **−4.76** | **−6.09 to −3.43** |
| $\hat{\theta}_{\text{simex}}^{(\text{cubic})}$ | 0.52 | 0.25 to 0.79 | −5.24 | −6.64 to −3.84 |

For heritability, the naive *h*^2^ estimator was found to be lower with respect to its actual value (Figure S3 of Appendix 1). A closer inspection of the additive 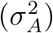, residual 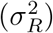 and permanent environmental 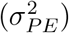 variance components indicated that this was due to an increase in 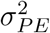 and a decrease in 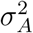.

For inbreeding depression, the error in the apparent pedigree induced an attenuation bias in *β_f_* that was comparable to case a) from the simulations (Figure 5b). The song sparrow pedigree has an architecture with *Ne/Nc* ≈ 0.6, which corresponds to a value between the simulated cases a) and b) with *Ne/Nc* = 1 and *Ne/Nc* = 0.3, respectively.

Interestingly, pedigree errors biased estimates of inbreeding depression more strongly than estimates of heritability (Table 1). This is not surprising, because information about inbreeding requires both parents to be known correctly, while information about additive genetic effects can be obtained from a single correctly assigned parent. As a consequence, pedigree error has more pronounced effects on estimates of inbreeding depression than on estimates of heritability. For both, inbreeding depression and heritability, different extrapolation functions yielded quite different estimates, which underlines the importance of the choice of the extrapolation function. This aspect follows from the approximate nature of the SIMEX method (Cook and Stefanski, 1994), and considerations about the choice of the extrapolant functions are reported in Carroll et al. (2006, chapter 5).

## Discussion

We have employed the case of pedigree error to promote a very general strategy to account for measurement error in ecology and evolution. By adapting the philosophy of the SIMEX approach, originally proposed to account for measurement error in continuous regression covariates (Cook and Stefanski, 1994), we illustrate how increasing error in the assignment of parents to their offspring in the pedigree can yield information about the resulting bias in parameters that are estimated in downstream statistical analyses, such as inbreeding depression or heritability. The observed trend with increasing error is then back-extrapolated to the hypothetical situation of zero error, yielding error-corrected, unbiased estimates of the parameters of interest. SIMEX is an intuitive way to assess and correct for the effects of any type of error in very general applications, especially when an explicit model for the unobserved component is difficult to formulate, and when it is thus not possible to embed an explicit error model directly in the statistical analysis. To facilitate the accessibility of the PSIMEX method, the code to perform all the analyses is provided via the novel R package `PSIMEX`, which is available from the CRAN repository (Ponzi, 2017).

We used simulation studies and a dataset from wild-living song sparrows to illustrate that the approach is successful in recovering error-corrected estimates of inbreeding depression and heritability in the presence of pedigree error. Interestingly, the simulations also revealed that pedigree errors do not necessarily lead to attenuated versions of quantitative genetic measures. In fact, inbreeding depression was over-*or* underestimated with increasing misassigned paternity error, and the direction of the bias depended on the pedigree topology. This result indicates that without observing or simulating the actual effect of the error, the direction of the bias in the naive estimator cannot be known a priori, even in the simple case of completely random error. Encouragingly, the 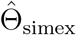 estimates obtained through the PSIMEX procedure captured these effects correctly and were always much closer to the true values than the naive estimates without error modelling. The application of the method to a wild population of song sparrows, where PSIMEX estimators could be compared to actual values derived from error-free pedigrees, confirmed that the method is performing well.

An important prerequisite to apply any error correction technique is knowledge of the error-generating mechanism and the error model parameter(s), as otherwise error models are nonidentifiable (Fuller, 1987; Gustafson, 2005; Carroll et al., 2006). Unlike in the case of other error-correction techniques, however, the SIMEX approach does not require that an explicit latent model for the unobserved, true covariate without measurement error is formulated. Moreover, we have illustrated in this paper that the specification of an error model for the variable of interest can be circumvented by knowing the error generating mechanism at the lowest level of the data generation process. In the case of PSIMEX, both the relatedness matrix **A** and the inbreeding coefficients *f_i_* of individuals are deduced from the pedigree, thus we did not have to formulate error models for **A** or *f_i_*, which would have been challenging (see, for example, Appendix 1, section 2.1). Instead, we could directly work with the error structure in the pedigree, which was much more straightforward. Of course, this required that information about the underlying error generating mechanism and the error proportion in the pedigree was available. In our application to the song sparrows, we knew the proportion of erroneously assigned fathers in the pedigree from a comparison of the error-prone and the error-free pedigree, and we assumed a random error mechanism where fathers are replaced with random individuals from the same generation, an assumption in line with available data (Reid et al., 2015).

To obtain unbiased, error-corrected estimates of parameters of interest using SIMEX, it is crucial to collect data that allows the quantification of the error. False assumptions may lead to biased SIMEX estimators, as illustrated in Appendix 1 (Sections 4 and 5). Therefore, it is extremely important to collect information about the error. Ideally, error estimation should be part of the study design, because it is much harder to obtain error estimates retrospectively. Quite often it will be sufficient to take error-prone and error-free measurements on a small subset of all study subjects. In the case of pedigrees, for example, it can be useful to genetically verify a subset of all parents in order to estimate the error proportion. Even in the absence of precise information about the error, similar studies or comparable populations might provide useful information, which can be used as *prior knowledge*. Of course, transportation of such information across study systems bears the risk that potentially inappropriate but untestable assumptions enter the modelling process, and it is therefore advisable to obtain some error estimates in the actual study system for comparison.

The PSIMEX methodology can be adapted to correct for different error mechanisms in the pedigree, for example when misassignments do not only affect fathers but also mothers, when the proportion of misassigned paternities varies across the study period, or when the replacement of fathers is more likely to occur with phenotypically or genotypically similar individuals. These error generating mechanisms can easily be included in the PSIMEX algorithm (see Section 5 of Appendix 1 for some examples). Moreover, the PSIMEX idea can be applied to virtually any quantity that is derived from pedigrees, for example to error-correct the estimates of variance parameters, but also to estimates of sexual selection, linkage, penetrance, the response to selection, genetic correlations, etc.

Although we have employed the SIMEX algorithm here only for a particular application to tackle pedigree error, the same generic principle can be adapted to many other situations. As an example, the SIMEX procedure could be used to account for location error in habitat selection studies, where parameters of interest, such as measures of distance and velocity, classifications of an animal’s activities, or the detection of an animal’s presence or absence at a given location may be biased (Ganskopp and Johnson, 2007; McKenzie et al., 2009). Instead of formulating an error model for the biased covariates themselves, it might often be easier to focus directly on the location error, using information on the accuracy of the measurements (*e. g*. GPS error) and the mechanisms that might obscure it, which can be used to obtain error model parameters for a SIMEX correction.

## Conclusions

The conceptual simplicity of the SIMEX philosophy allows its implementation even in situations when it is difficult or impossible to formulate or incorporate an explicit error model for an erroneous variable. The only prerequisites to apply the SIMEX algorithm are that the error-generating mechanism is known, and that it is possible to make the error “worse” in a controllable, quantitative way. We believe that many other applications in ecology and evolution might benefit from this simple and practical approach to obtain error-corrected parameter estimates in the presence of measurement error.

## Supporting information

**Appendix 1:** Supplementary text and figures (pdf)

**Appendix 2:** R script for PSIMEX on inbreeding in simulated data

**Appendix 3:** R script for PSIMEX on heritability in simulated data

**Appendix 4:** R script for PSIMEX on inbreeding

**Appendix 5:** R script for PSIMEX on heritability

**Appendix 6:** R script for PSIMEX on inbreeding in Song Sparrows

**Appendix 7:** R script for PSIMEX on heritability in Song Sparrows

## Author contributions

L.F.K. and S.M. conceived the research idea. E.P. designed and conducted the simulations and analyses. E.P. and S.M. wrote the manuscript. All authors provided feedback during the writing process and gave final approval.

## Acknowledgements

We thank Peter Arcese and Jane Reid for the use of the song sparrow data.

## REFERENCES

Apanasovich, T. V., R. J. Carroll, and A. Maity (2009). SIMEX and standard error estimation in semiparametric measurement error models. Electronic Journal of Statistics 3, 318–348.

Becker, P. J., J. Hegelbach, L. F. Keller, and E. Postma (2016). Phenotype-associated inbreeding biases estimates of inbreeding depression in a wild bird population. Journal of Evolutionary Biology 29, 35–46.

Bishop, D. A. and C. M. Beier (2013). Assessing uncertainty in high-resolution spatial climate data across the US Northeast. PLOS One 8, e70260.

Buonaccorsi, J. (2010). Measurement error: models, methods and applications. Boca Raton: CRC press.

Carroll, R. J., D. Ruppert, L. A. Stefanski, and C. M. Crainiceanu (2006). Measurement error in nonlinear models, a modern perspective. Boca Raton: Chapman and Hall.

Charlesworth, D. and J. H. Willis (2009). The genetics of inbreeding depression. Nature Reviews Genetics 10, 783–796.

Charmantier, A. and D. Reale (2005). How do misassigned paternities affect the estimation of heritability in the wild? Molecular Ecology 14, 2839–2850.

Cook, J. R. and L. A. Stefanski (1994). Simulation-extrapolation estimation in parametric measurement error models. Journal of the American Statistical Association 89, 1314–1328.

Coster, A. (2013). pedigree: Pedigree functions. R package version 1.4.

Devanarayan, V. and L. A. Stefanski (2002). Empirical simulation extrapolation for measurement error models with replicate measurements. Statistics and Probability Letters 59, 219–225.

Dohm, M. R. (2002). Repeatability estimates do not always set an upper limit to heritability. Functional Ecology 16, 273–280.

Eckert, R. S., R. J. Carroll, and N. Wang (1997). Transformations to additivity in measurement error models. Biometrics 53, 262–272.

Frankham, R. (1995). Effective population size/adult population size ratios in wildlife: a rewiev. Genetic Research 66, 95–107.

Freckleton, R. P. (2011). Dealing with collinearity in behavioural and ecological data: model averaging and the problems of measurement error. Behavioral Ecology and Sociobiology 65, 91–101.

Fuller, W. A. (1987). Measurement Error Models. New York: John Wiley & Sons.

Fung, K. Y. and D. Krewski (1999). On measurement error adjustment methods in Poisson regression. Environmetrics 10, 213–224.

Ganskopp, D. C. and D. D. Johnson (2007). GPS error in studies addressing animal movements and activities. Rangeland Ecology and Management 60, 350–358.

Ge, T., A. J. Holmes, R. L. Buckner, J. W. Smoller, and M. R. Sabuncu (2017). Heritability analysis with repeat measurements and its application to resting-state functional connectivity. PNAS 114, 5521–5526.

Germain, R. R., M. E. Wolak, P. Arcese, S. Losdat, and J. M. Reid (2016). Direct and indirect genetic and fine-scale location effects on breeding date in song sparrows. Journal of Animal Ecology 85, 1613–1624.

Gorjanic, G. and D. A. Henderson (2007). GeneticsPed: Pedigree and genetic relationship functions. R package version 1.34.0.

Gould, W. R., L. A. Stefanski, and K. H. Pollock (1999). Use of simulation-extrapolation estimation on catch-effort analysis. Canadian Journal of Fisheries and Aquatic Sciences 56, 1234–1240.

Griffith, S. C., I. P. Owens, and K. A. Thuman (2002). Extrapair paternity in birds: a review of interspecific variation and adaptive function. Molecular Ecology 11, 2195–2212.

Guelat, J. and M. Kery (2018). Effects of spatial autocorrelation and imperfect detection on species distribution models. Methods in Ecology and Evolution 9, 1614–1625.

Guillera-Arroita, G., J. J. Lahoz-Monfort, D. I. MacKenzie, B. A. Wintle, and M. A. McCarthy (2014). Ignoring imperfect detection in biological surveys is dangerous: a response to ‘Fitting and interpreting occupancy models’. PLOS One 9, 1–14.

Gustafson, P. (2004). Measurement Error and Misclassification in Statistics and Epidemiology: Impacts and Bayesian Adjustments. Boca Raton: Chapman & Hall/CRC.

Gustafson, P. (2005). On model expansion, model contraction, identifiability and prior information: Two illustrative scenarios involving mismeasured variables. Statistical Science 20, 111–140.

Hadfield, J. D. (2010). MCMC methods for multi-response generalized linear mixed models: The MCMCglmm R package. Journal of Statistical Software 33, 1–22.

Haila, Y., K. Henle, E. Apostolopoulou, J. Cent, E. Framstad, C. Goerg, K. Jax, R. Klenke, W. Magnuson, Y. G. Matsinos, B. Mueller, R. Paloniemi, J. Pantis, F. Rauschmayer, I. Ring, J. Settele, J. Simila, K. Touloumis, J. Tzanopoulos, and G. Pe’er (2014). Confronting and coping with uncertainty in biodiversity research and praxis. Nature Conservation 8, 45–75.

Henderson, C. R. (1976). Simple method for computing inverse of a numerator relationship matrix used in prediction of breeding values. Biometrics 32, 69–83.

Hoffmann, A. A. (2000). Laboratory and field heritabilities: Lessons from *Drosophila*. In T. Mousseau, S. B., and J. Endler (Eds.), Adaptive Genetic Variation in the Wild. New York, Oxford: Oxford Univ Press.

Holand, A., I. Steinsland, S. Martino, and H. Jensen (2013). Animal models and integrated nested Laplace approximations. G3 3, 1241–1251.

Hwang, W. H. and S. Y. H. Huang (2003). Estimation in capture-recapture models when covariates are subject to measurement errors. Biometrics 59, 1113–1122.

Jensen, H., E. M. Bremset, T. H. Ringsby, and B. E. Sæther (2007). Multilocus heterozygosity and inbreeding depression in an insular house sparrow metapopulation. Molecular Ecology 16, 4066–4078.

Keller, L. F., P. R. Grant, B. R. Grant, and K. Petren (2001). Heritability of morphological traits in Darwin’s Finches: misidentified paternity and maternal effects. Heredity 87, 325–336.

Keller, L. F., P. R. Grant, B. R. Grant, and K. Petren (2002). Environmental conditions affect the magnitude of inbreeding depression in survival of Darwin’s Finches. Evolution 56, 1229–1239.

Keller, L. F. and D. M. Waller (2002). Inbreeding effects in wild populations. Trends in Ecology and Evolution 17, 230–241.

Kruuk, L. E. B. (2004). Estimating genetic parameters in natural populations using the ‘animal model’. Philosophical Transactions of the Royal Society of London 359, 873–890.

Kruuk, L. E. B. and J. D. Hadfield (2007). How to separate genetic and environmental causes of similarity between relatives. Journal of Evolutionary Biology 20, 1890–1903.

Kuechenhoff, H., S. M. Mwalili, and E. Lesaffre (2006). A general method for dealing with misclassification in regression: The misclassification SIMEX. Biometrics 62, 85–96.

Lynch, M. and B. Walsh (1998). Genetics and Analysis of Quantitative Traits. Sunderland, MA: Sinauer Associates.

Macgregor, S., B. K. Cornes, N. G. Martin, and P. M. Visscher (2006). Bias, precision and heritability of self-reported and clinically measured height in Australian twins. Human Genetics 120, 571–580.

Mason, N. W. H., R. J. Holdaway, and S. J. Richardson (2018). Incorporating measurement error in testing for changes in biodiversity. Methods in Ecology and Evolution 9, 1296–1307.

McClintock, B. T., J. M. London, M. F. Cameron, and P. L. Boveng (2014). Modelling animal movement using the Argos satellite telemetry location error ellipse. Methods in Ecology and Evolution 6, 266–277.

McCullagh, P. and J. A. Nelder (1989). Generalized Linear models. London, UK: Chapman and Hall.

McKenzie, H. W., C. L. Jerde, D. R. Visscher, E. H. Merrill, and M. A. Lewis (2009). Inferring linear feature use in the presence of GPS measurement error. Environmental and Ecological Statistics 16, 531–546.

Melbourne, B. A. and P. Chesson (2006). The scale transition: scaling up population dynamics with field data. Ecology 87, 1478–1488.

Montgomery, R. A., G. J. Roloff, and J. M. Ver Hoef (2011). Implications of ignoring telemetry error on inference in wildlife resource use models. Journal of Wildlife Management 75, 702–708.

Muff, S. and L. F. Keller (2015). Reverse attenuation in interaction terms due to covariate error. Biometrical Journal 57, 1068–1083.

Muff, S., A. Riebler, L. Held, H. Rue, and P. Saner (2015). Bayesian analysis of measurement error models using integrated nested Laplace approximations. Journal of the Royal Statistical Society, Applied Statistics Series C 64, 231–252.

Palstra, F. P. and D. J. Fraser (2012). Effective/census population size ratio estimation: a compendium and appraisal. Ecology and Evolution 2, 2357–2365.

Ponzi, E. (2017). PSIMEX: SIMEX Algorithm on Pedigree Structures. University of Zurich. R package version 1.1.

Ponzi, E., L. F. Keller, T. Bonnet, and S. Muff (2018). Heritability, selection, and the response to selection in the presence of phenotypic measurement error: Effects, cures, and the role of repeated measurements. Evolution 0(0).

Reid, J. M., P. Arcese, and L. F. Keller (2006). Intrinsic parent-offspring correlation in inbreeding level in a song sparrow *(Melospiza melodia*) population open to immigration. The American Naturalist 168, 1–13.

Reid, J. M., P. Arcese, and L. F. Keller (2008). Individual phenotype, kinship, and the occurrence of inbreeding in song sparrows. Evolution 62, 887–899.

Reid, J. M., P. Arcese, L. F. Keller, R. R. Germain, A. B. Duthie, S. Losdat, M. E. Wolak, and P. Nietlisbach (2015). Quantifying inbreeding avoidance through extrapair reproduction. Evolution 69, 59–74.

Reid, J. M. and L. F. Keller (2010). Correlated inbreeding among relatives: occurrence, magnitude and implications. Evolution 64, 973–985.

Reid, J. M., L. F. Keller, A. B. Marr, P. Nietlisbach, R. J. Sardell, and P. Arcese (2014). Pedigree error due to extra-pair reproduction substantially biases estimates of inbreeding depression. Evolution 3, 802–815.

Rue, H., S. Martino, and N. Chopin (2009). Approximate Bayesian inference for latent Gaussian models by using integrated nested Laplace approximations (with discussion). Journal of the Royal Statistical Society. Series B (Statistical Methodology) 71, 319–392.

Sardell, R. J., L. F. Keller, P. Arcese, T. Bucher, and J. M. Reid (2010). Comprehensive paternity assignment: genotype, spatial location and social status in song sparrows *Melospiza melodia*. Molecular Ecology 19, 4352 – 4364.

Senneke, S. L., M. D. MacNeil, and L. D. Van Vleck (2004). Effects of sire misidentification on estimates of genetic parameters for birth and weaning weights in Hereford cattle. American Society of Animal Science 82, 2307–2312.

Smith, J. N. M. and L. F. Keller (2006). Genetics and demography of small populations. In J. Smith, L. Keller, A. Marr, and P. Arcese (Eds.), Conservation and biology of small populations: the song sparrows of Mandarte Island. Oxford University Press.

Smith, J. N. M. and R. Zach (1979). Heritability of some morphological characters in a song sparrow population. Evolution 33, 460–467.

Solow, A. R. (1998). On fitting a population model in the presence of observation error. Ecology 79, 1463–1466.

Stefanski, L. A. and J. R. Cook (1995). Simulation-extrapolation: The measurement error jackknife. Journal of the American Statistical Association 90, 1247–1256.

Steinsland, I., C. Larsen, A. Roulin, and H. Jensen (2014). Quantitative genetic modelling and inference in the presence of nonignorable missing data. Evolution 68, 1735–1747.

Stoklosa, J., C. Daly, S. D. Foster, M. B. Ashcroft, and D. I. Warton (2014). A climate of uncertainty: accounting for error in climate variables for species distribution models. Methods in Ecology and Evolution 6, 412–423.

van der Sluis, S., M. Verhage, D. Posthuma, and C. V. Dolan (2010). Phenotypic complexity, measurement bias, and poor phenotypic resolution contribute to the missing heritability problem in genetic association studies. PLOS One 5, e13929.

Visscher, P. M., J. A. Woolliams, J. Smith, and J. L. Williams (2002). Estimation of pedigree errors in the UK dairy population using microsatellite markers and the impact on selection. Journal of Dairy Science 85, 2368–2375.

Wilson, A. J., D. Reale, M. N. Clements, M. M. Morrissey, E. Postma, C. A. Walling, L. E. B. Kruuk, and D. H. Nussey (2010). An ecologist’s guide to the animal model. Journal of Animal Ecology 79, 13–26.

Wolak, M. E. and L. F. Keller (2014). Dominance genetic variance and inbreeding in natural populations. In A. Charmantier, D. Garant, and L. E. B. Kruuk (Eds.), Quantitative Genetics in the Wild. New York: Oxford University Press.

Wright, W. J., K. M. Irvine, J. M. Warren, and J. K. Barnett (2017). Statistical design and analysis for plant cover studies with multiple sources of observation errors. Methods in Ecology and Evolution 8, 1832–1841.

