## Supplementary material for "The simulation extrapolation technique meets ecology and evolution: A general and intuitive method to account for measurement error": R code: Appendix2.R

```
##############################################################
## The simulation extrapolation technique meets ecology and evolution: 
## A general and intuitive method to account for measurement error 
## R-code for P-SIMEX algorithm on inbreeding depression estimation:
## Simulation of pedigree and inbreeding depression data
## Generation of initial error on the simulated data
## Application of the P-SIMEX algorithm on the error prone data
################################################################


# install required libraries
source("https://bioconductor.org/biocLite.R")
biocLite("GeneticsPed")


install.packages('pedigree')
install.packages('pedigreemm')
install.packages('MCMCglmm')

# load required libraries
library("pedigree")
library("GeneticsPed") 
library("pedigreemm")
library("MCMCglmm")

# seed 
set.seed(49494)


# set a value for the Ne/Nc ratio
# here is 0.5 but it can be set to 0.15 or 0.05
NeNc <- 0.5
# set other parameters 
# (idgen = id per generations, nGen = number of generations)
idgen <- 100
nGen <- 30

# generate the pedigree using the function generatePedigree from GeneticsPed
# nFather and nMother are the number of fathers and mothers per generation
# they are generated according to the selected Ne/Nc ratio
ped0 <- generatePedigree(nId = idgen, nGeneration = nGen, 
                         nFather = idgen * NeNc, nMother = idgen * NeNc)

# Total number of individuals
Nid <- length(ped0$id)

# Vector with animal id
animal.id <- 1 : Nid

# calculate the inbreeding coefficient from the pedigree
# function calcInbreeding from the package 'pedigree'
# pedigree format for calcInbreeding
# must be: id - dam - sire
pedigree <- ped0[ , c('id','mother','father')]
names(pedigree) <- c("id", "dam", "sire")
# calculate inbreeding coefficients:
f_inb <- calcInbreeding(pedigree)

# inbreeding coefficient characteristics
mean(f_inb)
median(f_inb)
var(f_inb)
hist(f_inb)


# generate breeding values (to simulate trait) 
# using the function rbv from MCMCglmm
# fix the additive variance value, here is 0.3 but it can be any value
sigmaA <- 0.3
u <- rbv(pedigree, sigmaA) 

# simulate independent environmental effects
# fix the environmental variance value, here is 0.1 but it can be any value
sigmaE <- 0.1
e <- rnorm(Nid,0,sd=sqrt(sigmaE))

# fixed effects are mu, sex and inbreeding coefficient
# mu is the baseline, here is chosen as 10 but it can be any value
mu <- 10
sex <- ped0$sex


# simulate y
# we chose beta_sex = 2 and inbreeding depression = beta_f = -7
Y <- mu+2*sex-7*f_inb+u+e 

#generate dataset
trait.data <- data.frame(Y, sex, f_inb, animal.id)
trait.data$sex <- as.factor(sex)

# linear model to compute inbreeding depression
# it is done using a simple linear model 
# with the trait as a response 
# and the inbreeding coefficient as explanatory variable
# we also add sex as a covariate
# inbreeding depression is the regression coefficient of f_inb

model <- lm(Y ~ f_inb +sex,
            data  =  trait.data )

# inbreeding depression 
summary(model)$coefficients


#############################################
# generate some errors in the initial pedigree 
# exclude founders from this
founders <- ped0[which(is.na(ped0$father)), ]$id
founders_ped <- ped0[founders, c('id', 'father', 'mother') ]
pedigree_nofounders <- ped0[-founders, c('id', 'father', 'mother') ] #we'll generate errors here
# bring pedigrees into format for calcInbreeding, 
# which must be: animal - sire - dam
names(founders_ped) <- c("animal", "sire", "dam")
names(pedigree_nofounders) <- c("animal", "sire", "dam")

# How to add 10% error:
# choose 10% individuals from pedigree_nofounders and set their fathers as random
# random fathers are chosen from the male individuals of the previous generation
# excluding the real one

# choose 10% of the individuals (no founders)
lambda0 <- 0.1
m <- sample(1:length(pedigree_nofounders$animal), lambda0*length(pedigree_nofounders$animal))

# for each of the chosen individuals we substitute its father 
# with a random male from the previous generation
for (i in m) {
  # we identify the generation of the sample individual 
  # and select all males out of the previous generation
  generation <- ped0[which(ped0$id == pedigree_nofounders$animal[i]), ]$generation
  possible_fathers <- ped0[ped0$generation == generation-1 & ped0$sex == 1 ,  ]$id
  # we then exclude the real father (we want to replace it with someone else)
  possible_fathers <- possible_fathers[-which(possible_fathers == pedigree_nofounders[i, 'sire'])]
  # we sample one over the whole set of possible fathers 
  # and substitute the real father with it in the pedigree
  pedigree_nofounders$sire[i] <- possible_fathers[sample(1:length(possible_fathers), 1)]
  
}

# re calculate pedigree and inbreeding coefficient
# we need to merge the new pedigree with the founders 
# to calculate inbreeding coefficient on the whole pedigree
pedigree_sim <- rbind(founders_ped, pedigree_nofounders)
f_inb_err <- data.frame(pedigree_sim$animal, calcInbreeding(pedigree_sim))
names(f_inb_err) <- c("animal.id", "f_inb1")

# we include the new error-prone inbreeding coefficients in the dataset
trait.data <- merge(trait.data, f_inb_err, by = "animal.id") 
trait.data$f_err <- as.numeric(trait.data$f_inb1)

# compute initial values (error prone values)
model_err <-  lm(Y ~ f_err + sex, 
              data  =  trait.data )

# results
summary(model_err)$coefficients

# inbreeding coefficient characteristics
mean(trait.data$f_err)
median(trait.data$f_err)
var(trait.data$f_err)
hist(trait.data$f_err)

##############################
# implement PSIMEX on this data
# load PSIMEX library
library(PSIMEX)
# fixed error proportions
lambda <- c(0.2, 0.3, 0.4, 0.5, 0.6, 0.7, 0.8, 0.9, 1)
# initial error proportion
lambda0 <- 0.1

# initialize pedigree for PSIMEX
names(pedigree_sim) <- c('id', 'father', 'mother')
# include the information about generation and sex 
# from the initial simulated pedigree ped0
ped_simex <- merge(pedigree_sim, ped0, by = 'id')
# use just the error prone information and eliminate the true simulated one for PSIMEX
ped_simex <- ped_simex[,-c(2,3)]
names(ped_simex) <- c('id', 'father', 'mother', 'generation', 'sex')

# known dataset is the error prone one
trait_simex <- data.frame(trait.data$animal.id, trait.data$Y, trait.data$sex, trait.data$f_inb1)
names(trait_simex) <- c('id', 'Y', 'sex', 'f_inb')

# set model for PSIMEX
model_simex <- lm(Y ~ f_inb + sex, 
                  data  =  trait_simex )

# Psimex function
results.Psimex <- Psimex(ped_simex, data = trait_simex, lambda, lambda0, 
                  B = 100, model = model_simex, parameter = "inbreeding",
                  error = "misassignment", way = "uniform",
                  fitting.method = c("quadratic", "cubic"), 
                  ntop = NA, nbottom = NA, 
                  prior, nitt, thin, burnin)

# extract PSIMEX results
# initial information
results.Psimex$description
results.Psimex$error
results.Psimex$fitting.method
results.Psimex$way
results.Psimex$lambda
results.Psimex$lambda0 
results.Psimex$starting.value

# extrapolated values
results.Psimex$extrapolated_data        
```
