## Supplementary material for "The simulation extrapolation technique meets ecology and evolution: A general and intuitive method to account for measurement error": R code: Appendix3.R

```
##############################################################
## The simulation extrapolation technique meets ecology and evolution: 
## A general and intuitive method to account for measurement error 
## R-code for P-SIMEX algorithm on heritability estimation
## Simulation of pedigree and heritability data
## Generation of initial error on the simulated data
## Application of the P-SIMEX algorithm on the error prone data
##############################################################


# install required libraries
source("https://bioconductor.org/biocLite.R")
biocLite("GeneticsPed")


install.packages('pedigree')
install.packages('pedigreemm')
install.packages('MCMCglmm')

# load required libraries
library("pedigree")
library("GeneticsPed") 
library("pedigreemm")
library("MCMCglmm")

# seed 
set.seed(49494)


# set a value for the Ne/Nc ratio
# here is 0.5 but it can be set to 0.15 or 0.05
NeNc <- 0.5
# set other parameters (id per generations, number of generations)
idgen <- 100
nGen <- 30

# generate the pedigree using the function generatePedigree from GeneticsPed
# nFather and nMother are the number of fathers and mothers per generation
# they are generated according to the selected Ne/Nc ratio
ped0 <- generatePedigree(nId = idgen, nGeneration = nGen, 
                         nFather = idgen * NeNc, nMother = idgen * NeNc)

# Total number of individuals
Nid <- length(ped0$id)

# Vector with animal id
id <- 1 : Nid

# bring pedigree into format for rbv, which must be: id - mother - father
pedigree <- ped0[ , c('id','mother','father')]
names(pedigree) <- c("id", "dam", "sire")

# generate breeding values using the function rbv from MCMCglmm
# fix the additive variance value, here is 0.3 but it can be any value
sigmaA <- 0.3
u <- rbv(pedigree, sigmaA) 

# simulate independent environmental effects
# fix the environmental variance value, here is 0.1 but it can be any value
sigmaE <- 0.1
e <- rnorm(Nid,0,sd=sqrt(sigmaE))

# fixed effects are mu and sex
# mu is the baseline, here is chosen as 10 but it can be any value
mu <- 10
sex <- ped0$sex

# simulate y
# we chose beta_sex = 2 
Y <- mu+2*sex+u+e

# generate dataset
trait.data <- data.frame(Y, sex, id)
trait.data$sex <- as.factor(sex)
trait.data <- merge(trait.data, ped0[, c('id', 'generation')], by = 'id')
# year is the generation number
names(trait.data) <- c('animal', 'Y', 'sex', 'year')


# MCMCglmm to compute heritability (VA/(VA+VE))
# (it should be 0.3/(0.3+0.1) = 0.75)
# it decomposes the variance into additive variance (VA, given by animal id)
# and environmental variance (VE, as residual variance)
# one could add more random effects if more components are needed

# order pedigree to fulfill MCMCglmm requirements
ord <- orderPed(pedigree)
pedigree <- pedigree[order(ord),]

# priors are set as inverse gammas
# the prior assumption is that the phenotypic variance 
# is equally divided into the two components 
p.var <- var(trait.data$Y,na.rm = TRUE)
prior <- list(G = list(G1 = list(V = matrix(p.var/2), n = 1)),
              R = list(V = matrix(p.var/2),n=1))

# MCMCglmm model has sex as fixed effect 
# and the animal id as random effect (which gives VA)

model <- MCMCglmm(Y ~ sex, random = ~animal,
                  pedigree = pedigree, data = trait.data, prior = prior,
                  nitt = 11000, thin = 100, burnin = 1000, verbose = F)

# extract heritability: VA/(VA+VE)
# if other random effects are present they have to be added in the denominator
heritability <- model$VCV[,"animal"]/
  (model$VCV[,"animal"]+model$VCV[,"units"])

# mean value for heritability
mean(heritability) 
# standard error for heritability
sd(heritability)
# additive component of variance
mean(model$VCV[,"animal"])
# environmental component of variance
mean(model$VCV[,"units"])

#############################################
# generate some errors in the initial pedigree 
# exclude founders from this
founders <- ped0[which(is.na(ped0$father)), 1]
founders_ped <- ped0[founders, 1:3 ]
pedigree_nofounders <- ped0[-founders, 1:3 ] #we'll generate errors here
# bring pedigrees into format for MCMCglmm, which must be: id - mother - father
names(founders_ped) <- c("animal", "sire", "dam")
names(pedigree_nofounders) <- c("animal", "sire", "dam")

# How to add 10% error:
# choose 10% individuals from pedigree_nofounders and set their fathers as random
# random fathers are chosen from the male individuals of the previous generation
# excluding the real one

# choose 10% of the individuals (no founders)
lambda0 <- 0.1
m <- sample(1:length(pedigree_nofounders$animal), lambda0*length(pedigree_nofounders$animal))

# for each of the chosen individuals we substitute its father 
# with a random male from the previous generation
for (i in m) {
  # we identify the generation of the sample individual 
  # and select all males out of the previous generation
  generation <- ped0[which(ped0$id == pedigree_nofounders$animal[i]), ]$generation
  possible_fathers <- ped0[ped0$generation == generation-1 & ped0$sex == 1 ,  ]$id
  # we then exclude the real father (we want to replace it with someone else)
  possible_fathers <- possible_fathers[-which(possible_fathers == pedigree_nofounders[i, 'sire'])]
  # we sample one over the whole set of possible fathers 
  # and substitute the real father with it in the pedigree
  pedigree_nofounders$sire[i] <- possible_fathers[sample(1:length(possible_fathers), 1)]
  
}

# re calculate pedigree 
# we need to merge the new pedigree with the founders 
# to calculate variance components on the whole pedigree
pedigree_err <- rbind(founders_ped, pedigree_nofounders)
# order pedigree to fulfill MCMCglmm requirements
ord <- orderPed(pedigree_err)
pedigree_err <- pedigree_err[order(ord),]
# compute initial values (error prone values)
# we use the error-prone pedigree in the MCMCglmm model
model_err <- MCMCglmm(Y ~ sex, random = ~animal,
                   pedigree = pedigree_err, data = trait.data, prior = prior,
                   nitt = 11000, thin = 100, burnin = 1000, verbose = F)

# extract heritability: VA/(VA+VE)
# use the same formula as in the first model
heritability_err <- model_err$VCV[,"animal"]/
  (model_err$VCV[,"animal"]+model_err$VCV[,"units"])

# mean value for heritability
mean(heritability_err) 
# standard error for heritability
sd(heritability_err)
# additive component of variance
mean(model_err$VCV[,"animal"])
# environmental component of variance
mean(model_err$VCV[,"units"])

##############################
# implement SIMEX on this data
# load PSIMEX library
library(PSIMEX)

# fixed error proportions
lambda <- c(0.2, 0.3, 0.4, 0.5, 0.6)
# initial error proportion
lambda0 <- 0.1

# initialize pedigree for PSIMEX
names(pedigree_err) <- c('id', 'father', 'mother')
# include the information about generation and sex 
# from the initial simulated pedigree ped0
ped_simex0 <- merge(pedigree_err, ped0, by = 'id')
# use just the wrong information and eliminate the true simulated one for PSIMEX
ped_simex0 <- ped_simex0[,-c(2,3)]
names(ped_simex0) <- c('id', 'father', 'mother', 'generation', 'sex')


# to fulfill MCMCglmm requirements
ped_simex <- ped_simex0[ , c('id','father','mother')]
names(ped_simex) <- c("animal", "sire", "dam")
ord <- orderPed(ped_simex)
ped_simex <- ped_simex[order(ord),]


# set MCMCglmm for PSIMEX
model <- MCMCglmm(Y~1+sex, random = ~animal,
                  pedigree = ped_simex, data = trait.data,
                  prior = prior, nitt = 11000, thin = 100,
                  burnin = 1000,
                  verbose = FALSE)

#to fulfill PSIMEX requirements 
trait.data$id <- trait.data$animal

# Psimex function
results.Psimex <- Psimex(ped_simex0, data = trait.data, lambda, lambda0, 
                         B = 100, model, parameter = "heritability",
                         error = "misassignment", way = "uniform",
                         fitting.method = "line",
                         ntop = NA, nbottom = NA,
                         prior = prior, nitt = 11000, thin = 100,
                         burnin = 1000)

# extract PSIMEX results
# initial information
results.Psimex$description
results.Psimex$error
results.Psimex$fitting.method
results.Psimex$way
results.Psimex$lambda
results.Psimex$lambda0 
results.Psimex$starting.value

# extrapolated values
results.Psimex$extrapolated_data        
```
