## Supplementary material for "The simulation extrapolation technique meets ecology and evolution: A general and intuitive method to account for measurement error": R code: Appendix4.R

```
##############################################################
## The simulation extrapolation technique meets ecology and evolution: 
## A general and intuitive method to account for measurement error 
## R-code for P-SIMEX algorithm on inbreeding depression estimation
## Application of P-SIMEX on an error prone pedigree
##############################################################


# load PSIMEX library
library("PSIMEX")

#load datasets for pedigree and trait data
pedigree0 <- read.table('pedigree.txt')
data <- read.table('trait.txt')

head(pedigree0)
head(data)

# initial error proportion is assumed to be 10%
lambda0 <- 0.1

# fix increasing error proportion for PSIMEX
lambda <- c(0.2, 0.3, 0.4, 0.5, 0.6)

# linear model to compute inbreeding depression
# it is done using a simple linear model 
# with the trait as a response 
# and the inbreeding coefficient as explanatory variable
# we also add sex as a covariate
# inbreeding depression is the regression coefficient of f_inb
model <- lm(y ~ sex+f_inb, data = data)


# Psimex function for misassignement error 
# uniformly distributed across the pedigree
# we set a number of simulations equal to 100 
# and a quadratic extrapolation function
results.Psimex <- Psimex(pedigree0, data, lambda, lambda0, 
                         B = 100, model, parameter = "inbreeding", 
                         error = "misassignment", way = "uniform", 
                         fitting.method = "quadratic")

# extract PSIMEX results
# initial information
results.Psimex$description
results.Psimex$error
results.Psimex$fitting.method
results.Psimex$way
results.Psimex$lambda
results.Psimex$lambda0 
results.Psimex$starting.value

# extrapolated values
results.Psimex$extrapolated_data 
```
