## Supplementary material for "The simulation extrapolation technique meets ecology and evolution: A general and intuitive method to account for measurement error": R code: Appendix5.R

```
##############################################################
## The simulation extrapolation technique meets ecology and evolution: 
## A general and intuitive method to account for measurement error 
##
## R-code for P-SIMEX algorithm on heritability estimation
## Application of P-SIMEX on an error prone pedigree
##############################################################

# load PSIMEX library
library("PSIMEX")

#load datasets for pedigree and trait data
pedigree0 <- read.table('pedigree.txt')
data <- read.table('trait.txt')

head(pedigree0)
head(data)

# initial error proportion is assumed to be 10%
lambda0 <- 0.1

# fix increasing error proportion for PSIMEX
lambda <- c(0.2, 0.3, 0.4, 0.5, 0.6)

# prior are set as inverse gammas
# variance is assumed to be equally divided into the three components:
# VA = additive genetic variance
# VPE = permanent environmental variance
# VE = environmental variance
# h = VA/ (VA+VPE+VE)
prior <- list(G=list(G1=list(V=matrix(1/3),n=1),
                     G2=list(V=matrix(1/3),n=1)),
              R=list(V=matrix(1/3),n=1))

#to fulfill MCMCglmm requirements
pedigree <- pedigree0[ , c(1,2,3)]
names(pedigree) <- c("animal", "dam", "sire")
ord <- orderPed(pedigree)
pedigree <- pedigree[order(ord),]


# MCMCglmm model has sex and baseline as fixed effect 
# and the animal id as random effect (which gives VA)
# plus an additional id as random effect to model permanent environment effect (VPE)
model <- MCMCglmm(y ~ 1+sex, random = ~animal+id, 
                  pedigree = pedigree, data = data, 
                  prior = prior, nitt = 11000, thin = 100, burnin = 1000, 
                  verbose = FALSE)


# Psimex function for misassignement error uniformly distributed across the pedigree
# we set a number of simulations equal to 10 and a quadratic extrapolation 
# we need to set parameters for MCMCglmm as well (nitt, burnin, thin)
results.Psimex <- Psimex(pedigree0, data, lambda, lambda0, 
                  B = 100, model, parameter = "heritability", 
                  error = "misassignment", way = "uniform", 
                  fitting.method = c("linear", "quadratic"), 
                  ntop = NA, nbottom = NA, 
                  prior = prior, nitt = 11000, thin = 100, burnin = 1000)

# extract PSIMEX results
# initial information
results.Psimex$description
results.Psimex$error
results.Psimex$fitting.method
results.Psimex$way
results.Psimex$lambda
results.Psimex$lambda0 
results.Psimex$starting.value

# extrapolated values
results.Psimex$extrapolated_data 
```
