## Supplementary material for "The simulation extrapolation technique meets ecology and evolution: A general and intuitive method to account for measurement error": R code: Appendix6.R

```
##############################################################
## The simulation extrapolation technique meets ecology and evolution: 
## A general and intuitive method to account for measurement error  
##
## R-code for P-SIMEX algorithm on inbreeding depression
## in song sparrows data
##############################################################


# control function
is.integer0 <- function(x)
{
  is.integer(x) && length(x) == 0L
}


library(pedigree)
library(MCMCglmm)
library(parallel)
library(lme4)

#Load datasets 
allbirds <- read.csv("SongSparrowData_noembr.csv", sep="\t")
allbirds$ninecode <- allbirds$id
# calculate life time and survival
allbirds$life <- allbirds$death.year- allbirds$birth.year
# juvenile survival is 1 if life is at least one year, 0 otherwise
allbirds$js <- as.numeric(allbirds$life>0)

# calculate generation as birth year
allbirds$generation_dad <- c()
for (jj in 1:length(allbirds[ ,1])){
  
  mm <- which(allbirds$id == allbirds$social.father[jj])
  if (is.integer0(mm)) allbirds$generation_dad[jj] <- NA
  if (!is.integer0(mm)) allbirds$generation_dad[jj] <- allbirds[mm, ]$birth.year
  
  
}

allbirds <- allbirds[which(!is.na(allbirds$generation_dad)), ]
allbirds <- allbirds[which(!duplicated(allbirds$ninecode)), ]


# true value to compare
# from genetic pedigree
# use f.corr from data set
# store covariates of interest
trait1 <- data.frame(allbirds$id, allbirds$js, allbirds$f.corr,  allbirds$sex, allbirds$Year)
names(trait1) <- c("ID", "h", "f", "sex", "Year")


# calculate true estimate of inbreeding depression
# glm model with cloglog link
# sex and year as additional covariates

model1 <- glm(h ~ f+sex+Year,
              data  =  trait1, 
              family = binomial(link = cloglog))

# extract inbreeding depression and its SE
true <- summary(model1)$coefficients[2,1]
se_true <- summary(model1)$coefficients[2,2]


# extract social pedigree 
pedigree <- data.frame(allbirds$ninecode, allbirds$social.mother, allbirds$social.father) 
names(pedigree) <- c("id", "dam", "sire")

# order pedigree
ord <- orderPed(pedigree)
pedigree <- pedigree[order(ord), ]
# calculate inbreeding coefficient from pedigree
f_inb <- data.frame(pedigree$id, calcInbreeding(pedigree))
names(f_inb) <- c("ID", "f_sim")

# store covariates of interest
trait <- data.frame(allbirds$id, allbirds$js)
names(trait) <- c("ID", "h")
trait <- merge(trait, f_inb, by = "ID")
trait$f <- as.numeric(trait$f_sim)
trait <- data.frame(allbirds$ninecode, allbirds$js, trait$f, allbirds$sex, allbirds$Year)
names(trait) <- c("ID", "h", "f", "sex", "Year")

# calculate naive estimate of inbreeding depression
# glm model with cloglog link
# sex and year as additional covariates

model <- glm(h ~ f+sex+Year, 
            data  =  trait, 
            family = binomial(link = cloglog))

# extract inbreeding depression and its SE
inb0 <- summary(model)$coefficients[2,1]
se_inb0 <-summary(model)$coefficients[2,2]
pval0 <- summary(model)$coefficients[2,4]

# store inbreeding coefficient characteristics
mean_inb0 <- mean(trait$f)
median_inb0 <- median(trait$f)
var_inb0 <- var(trait$f)


# apply PSIMEX
# increasing error proportions 
lambda0 <- c(0.2, 0.3, 0.4, 0.5,  0.6, 0.7, 0.8)
# actual error proportion needed 
lambda <- 1- (1-lambda0)/(1-0.17) 
# total number of PSIMEX iteration
nsim <- 100

# check if parentages are all known
known <- pedigree[which(!is.na(pedigree$dam)), ]
unknown <- pedigree[which(is.na(pedigree$dam)), ]


#setting initial values
inb <- matrix(data  =  NA, nrow  =  length(lambda), ncol  =  nsim)
se_inb <- matrix(data  =  NA, nrow  =  length(lambda), ncol  =  nsim)
mean_inb <- matrix(data  =  NA, nrow  =  length(lambda), ncol = nsim)
median_inb <- matrix(data  =  NA, nrow  =  length(lambda), ncol = nsim)
var_inb <- matrix(data  =  NA, nrow  =  length(lambda), ncol = nsim)

# repeat for each error proportions
for (k in 1:length(lambda)){
  # repeat for nsim simulations
  for (j in 1:nsim) {
    animal <- known$id
    dam <- known$dam
    sire <- known$sire  
    #initializing inbreeding coefficient
    trait$f_sim <- c()
    
    # choose a proportion of lambda over the total number of individuals 
    # and set their parents as unknown
    m <- sample(1:length(known$id), lambda[k]*length(known$id))
    for (i in m) {
      # sample possible replacements from the sam generation
      generation <- allbirds[which(allbirds$id == known$id[i]), ]$generation_dad
      # exclude the real father
      others <- allbirds[which(allbirds$birth.year == generation & allbirds$sex == "Male"),  ]$id
      # check if there are other possible individuals and pick one to replace 
      if (!is.integer0(others)){
        if (is.element(known[i, 3], others))  others <- others[-which(others == known[i, 3])]
        sire[i] <- others[sample(1:length(others), 1)]
      } 
    }
    
    #recalculate pedigree
    ped <- data.frame(id = animal, dam = dam, sire = sire)
    ped <- rbind(ped, unknown)
    ord <- orderPed(ped)
    ped <- ped[order(ord),]
    
    #re calculate inbreeding coefficient
    f_sim <- data.frame(ped$id, calcInbreeding(ped))
    names(f_sim) <- c("ID", "f_sim")
    # include it in the data
    trait <- merge(trait, f_sim, by = "ID")
    trait$f_sim <- as.numeric(trait$f_sim)
    
    # calculate inbreeding depression 
    if(sum(trait$f_sim)>0) {
      
      # glm for survival 
      # cloglog link function 
      # sex and year as covariates
      model <- glm(h ~ f_sim+sex+Year , 
                   data  =  trait, 
                   family = binomial(link = cloglog))
      
      # extract inbreeding depression and its SE
      inb[k, j] <- summary(model)$coefficients[2,1]
      se_inb[k,j] <- summary(model)$coefficients[2,2]
      
      # store inbreeding coefficient characteristics
      mean_inb[k, j] <- mean(trait$f_sim)
      median_inb[k, j] <- median(trait$f_sim)
      var_inb[k, j] <- var(trait$f_sim)
      
    }
    
    #this must be checked
    if(sum(trait$f_sim) == 0) {
      inb[k, j] <- NA
      se_inb[k,j] <- NA
      mean_inb[k, j] <- NA
      median_inb[k, j] <- NA
      var_inb[k, j] <- NA
    }
    
    
  }
  
}

#compute means across simulations
inb_ave <- rowMeans(inb, na.rm = TRUE)
se_inb_ave <- rowMeans(se_inb, na.rm = TRUE)
mean_inb_ave <- rowMeans(mean_inb, na.rm = TRUE)
median_inb_ave <- rowMeans(median_inb, na.rm = TRUE)
var_inb_ave <- rowMeans(var_inb, na.rm = TRUE)

# merge all values with naive ones
lambda <- c(0.17,lambda0)
inb_ave <- c(inb0, inb_ave)
se_inb <- c(se_inb0, se_inb_ave)
mean_inb <- c(mean_inb0,mean_inb_ave)
median_inb <- c(median_inb0, median_inb_ave)
var_inb <- c(var_inb0, var_inb_ave)

low <- inb_ave-1.96*se_inb
up <- inb_ave+1.96*se_inb


# Extrapolation phase of PSIMEX
p.names <- c("inb", "se_inb")
estimates <- data.frame(inb_ave, se_inb)
colnames(estimates) <- p.names

#linear case
extrapolation_inb <- lm(estimates[ ,1] ~ lambda)
inb_pred1 <- predict(extrapolation_inb, newdata = data.frame(lambda = 0))
extrapolation_inb_se <- lm(estimates[,2] ~ lambda)
inb_pred_se1 <- predict(extrapolation_inb_se, newdata = data.frame(lambda = 0))

#quadratic case
extrapolation_inb1 <- lm(estimates[ ,1] ~ lambda+ I(lambda^2))
inb_pred2 <- predict(extrapolation_inb1, newdata = data.frame(lambda = 0))
extrapolation_inb1_se <- lm(estimates[ ,2] ~ lambda+ I(lambda^2))
inb_pred1_se2 <- predict(extrapolation_inb1_se, newdata = data.frame(lambda = 0))

#cubic case
extrapolation_inb2 <- lm(estimates[ ,1] ~ lambda+ I(lambda^2)+ I(lambda^3))
inb_pred3 <- predict(extrapolation_inb2, newdata = data.frame(lambda = 0))
extrapolation_inb2_se <- lm(estimates[ ,2] ~ lambda+ I(lambda^2)+ I(lambda^3))
inb_pred1_se3 <- predict(extrapolation_inb2_se, newdata = data.frame(lambda = 0))


# calculate SIMEX SE

# first component 
# sampling variability
S <- c()
#one per lambda 
for ( i in 1:length(lambda0)) {
  diff <- c()
  #calculate the differences per each simulation
  for ( j in 1: nsim ) {
    diff[j] <- inb[i,j]-inb_ave[i]
  }
  
  diff <- na.omit(diff)
  S[i] <- 1/(nsim-1)*sum(diff^2) 
  
}
S <- c(se_inb_ave[1], S)

# extrapolated value
# linear case
extrapolation_var<- lm(S ~ lambda)
var_pred<-predict(extrapolation_var, newdata = data.frame(lambda = 0))

#quadratic case
extrapolation_var1<- lm(S ~ lambda+ I(lambda^2))
var_pred1<-predict(extrapolation_var1, newdata = data.frame(lambda = 0))

#cubic case
extrapolation_var2<- lm(S ~ lambda+ I(lambda^2)+ I(lambda^3))
var_pred2<-predict(extrapolation_var2, newdata = data.frame(lambda = 0))

# TOTAL SE
#linear 
var_ext1 <- inb_pred_se1^2 + var_pred

#quadratic
var_ext2 <- inb_pred1_se2^2 + var_pred1

#cubic
var_ext3 <- inb_pred1_se3^2 + var_pred2

true_low <- true - 1.96*se_true
true_up <- true + 1.96*se_true

naive <- inb0
naive_low <- naive - 1.96*se_inb0
naive_up <- naive + 1.96*se_inb0

inb_pred_low1 <- inb_pred1 - 1.96*sqrt(var_ext1)
inb_pred_up1 <- inb_pred1 + 1.96*sqrt(var_ext1)

inb_pred_low2 <- inb_pred2 - 1.96*sqrt(var_ext1)
inb_pred_up2 <- inb_pred2 + 1.96*sqrt(var_ext1)

inb_pred_low3 <- inb_pred3 - 1.96*sqrt(var_ext1)
inb_pred_up3 <- inb_pred3 + 1.96*sqrt(var_ext1)

AIC(extrapolation_inb)

AIC(extrapolation_inb2)

AIC(extrapolation_inb1)

results_sparrows_inb <- list(true, true_low, true_up, 
                             naive, naive_low, naive_up, 
                             inb_pred1, inb_pred_low1, inb_pred_up1, 
                             inb_pred2, inb_pred_low2, inb_pred_up2, 
                             inb_pred3, inb_pred_low3, inb_pred_up3)

save(results_sparrows_inb, file = 'results_sparrows_inb.RData')
```
