## Supplementary material for "The simulation extrapolation technique meets ecology and evolution: A general and intuitive method to account for measurement error": R code: Appendix7.R

```
##############################################################
## The simulation extrapolation technique meets ecology and evolution: 
## A general and intuitive method to account for measurement error 
##
## R-code for P-SIMEX algorithm on heritability
## in song sparrows data
##############################################################
## R version 3.3.2


library(synbreed)
library(pedigree)
library(MCMCglmm)

# control function
is.integer0 <- function(x)
{
  is.integer(x) && length(x) == 0L
}


  
 

#Load datasets 
allbirds<-read.csv("SongSparrowData_noembr.csv", sep="\t")
morpho<-read.csv("MandarteBiosBasicData2007.csv")

w<-intersect(allbirds$id, morpho$ninecode)
k<-which(is.element(morpho$ninecode, w))

morpho<-morpho[k, ]

morpho$sex<-c()
for ( i in 1:length(morpho[ ,1])) {
  w<-which(allbirds$id==morpho$ninecode[i])
  morpho$sex[i]<-allbirds$sex[w]
  
  
}

dati<-morpho[!is.na(morpho$sex), ]


# calculate true value
# from genetic pedigree
all.ids <- c(allbirds$id, allbirds$gen.father, allbirds$gen.mother)
all.ids <- all.ids[!duplicated(all.ids)]
all <- which(!is.na(all.ids))
gen.dad <- c()
gen.mom <- c()
idg <- c()

for (i in all) {
  w <- which(allbirds$id == all.ids[i])
  idg[i] <- allbirds$id[w]
  gen.dad[i] <- allbirds$father[w]
  gen.mom[i] <- allbirds$mother[w]
  
}

# store genetic pedigree
gen <- data.frame(id = idg, dam = gen.mom, sire = gen.dad)
# order pedigree
ord <- orderPed(gen)
gen <- gen[order(ord),]
# calculate inbreeding coefficient
f <- data.frame(gen$id, calcInbreeding(gen))
names(f) <- c("ID", "f")

# store variables of interest
pheno <- data.frame(dati$ninecode, dati$mintars, dati$sex)
names(pheno) <- c("animal", "tarsus", "sex")
pheno$ID <- pheno$animal
pheno <- merge(pheno, f, by = "ID")

# prior for MCMCglmm
# inverse gamma priors for all components
# all the same proportion of variance
p.var <- var(pheno$tarsus, na.rm = TRUE)
prior <- list(G = list(G1 = list(V = matrix(p.var/3), n = 1),
                       G2 = list(V = matrix(p.var/3), n = 1)),
              R = list(V = matrix(p.var/3), n = 1))

# MCMCglmm for tarsus length
# fixed effects are sex and inbreeding
# random effects are the animal (VA)
# and the ID (VPE)
model <- MCMCglmm(tarsus ~ 1+sex+f, random = ~animal+ID,
                pedigree = gen, data = pheno, prior = prior,
                nitt = 11000, thin = 100, burnin = 1000, verbose = T)

# extract results from MCMCglmm
posterior.heritability <- model$VCV[,"animal"]/
  (model$VCV[,"animal"]+model$VCV[,"units"]+model$VCV[,"ID"])

h_true <- mean(posterior.heritability) 
se_true <- sd(posterior.heritability) 
VA_true<-mean(model$VCV[,"animal"])
VE_true<-mean(model$VCV[,"units"])
VPE_true<-mean(model$VCV[,"ID"])


# calculate naive value
# from social pedigree 
all.ids1 <- c(allbirds$id, allbirds$social.father, allbirds$social.mother)
all.ids1 <- all.ids1[!duplicated(all.ids1)]
all1 <- which(!is.na(all.ids1))
soc.dad <- c()
soc.mom <- c()
idg <- c()

for (i in all1) {
  w<-which(allbirds$id==all.ids1[i])
  idg[i]<-allbirds$id[w]
  soc.dad[i]<-allbirds$social.father[w]
  soc.mom[i]<-allbirds$social.mother[w]
  
}

# store social pedigree
soc <- data.frame(id = idg, dam = soc.mom, sire = soc.dad)
# order pedigree
ord <- orderPed(soc)
soc <- soc[order(ord),]

# calculate inbreeding coefficients
f <- data.frame(soc$id, calcInbreeding(soc))
names(f) <- c("ID", "f")

# store variables of interest
pheno <- data.frame(dati$ninecode, dati$mintars, dati$sex)
names(pheno) <- c("animal", "tarsus", "sex")
pheno$ID <- pheno$animal
pheno <- merge(pheno, f, by = "ID")


# MCMCglmm for tarsus length
# fixed effects are sex and inbreeding
# random effects are the animal (VA)
# and the ID (VPE)
model1 <- MCMCglmm(tarsus ~ 1+sex+f, random = ~animal+ID,
                pedigree = soc, data = pheno, prior = prior,
                nitt = 11000, thin = 100, burnin = 1000, verbose = T)

posterior.heritability <- model1$VCV[,"animal"]/
  (model1$VCV[,"animal"]+model1$VCV[,"units"]+model1$VCV[,"ID"])

h0 <- mean(posterior.heritability) 
s0 <- sd(posterior.heritability) 
VA0 <- mean(model1$VCV[,"animal"])
VE0 <- mean(model1$VCV[,"units"])
VPE0 <- mean(model1$VCV[,"ID"])

allbirds$generation_dad <- c()
for (jj in 1:length(allbirds[ ,1])){
  
  mm <- which(allbirds$id == allbirds$social.father[jj])
  if (is.integer0(mm)) allbirds$generation_dad[jj] <- NA
  if (!is.integer0(mm)) allbirds$generation_dad[jj] <- allbirds[mm, ]$birth.year
  
  
}


# apply PSIMEX
# increasing error proportions 
lambda0<-c(0.2, 0.3, 0.4, 0.5,  0.6, 0.7)
# actual error proportion needed 
lambda <- 1- (1-lambda0)/(1-0.17)
# total number of PSIMEX iteration

nsim<-100

# start with the social parents
all.ids<-c(allbirds$id, allbirds$social.father, allbirds$social.mother)
all.ids<-all.ids[!duplicated(all.ids)]
all<-which(!is.na(all.ids))


gen.dad<-c()
gen.mom<-c()
idg<-c()

for (i in all) {
  w<-which(allbirds$id==all.ids[i])
  idg[i]<-allbirds$id[w]
  gen.dad[i]<-allbirds$social.father[w]
  gen.mom[i]<-allbirds$social.mother[w]
  
}

# Simulation phase of PSIMEX
# setting initial values
h <- matrix(data = NA, nrow = length(lambda), ncol = nsim)
se_h <- matrix(data = NA, nrow = length(lambda), ncol = nsim)
VA <- matrix(data = NA, nrow = length(lambda), ncol = nsim)
VE <- matrix(data = NA, nrow = length(lambda), ncol = nsim)
VPE <- matrix(data = NA, nrow = length(lambda), ncol = nsim)


# repeat for each error proportions
for (k in 1:length(lambda)){
  # repeat for nsim simulations
  for (j in 1:nsim) {
    
    
    # choose a proportion of lambda over the total number of individuals 
    # and set their parents as unknown
    m <- sample(1:length(allbirds$id), lambda[k]*length(allbirds$id))
    for (i in m) {
      # sample possible replacements from the sam generation
      generation <- allbirds[i, ]$generation_dad
      # exclude the real father
      others <- allbirds[which(allbirds$birth.year == generation & allbirds$sex == "Male"),  ]$id
      # check if there are other possible individuals and pick one to replace 
      if (!is.integer0(others)){
        if (is.element(gen.dad[i], others))  others <- others[-which(others == gen.dad[i])]
        gen.dad[i] <- others[sample(1:length(others), 1)]
        
      }
    }
    
    #recalculate pedigree
    gen <- data.frame(id=idg, dam=gen.mom, sire=gen.dad)
    ord <- orderPed(gen)
    gen <- gen[order(ord),]
    
    #re calculate inbreeding coefficient
    f <- data.frame(gen$id, calcInbreeding(gen))
    names(f) <- c("ID", "f")
    
    # store variables of interest
    pheno <- data.frame(dati$ninecode, dati$mintars, dati$sex)
    names(pheno) <- c("animal", "tarsus", "sex")
    pheno$ID <- pheno$animal
    pheno <- merge(pheno, f, by = "ID")
    
    # MCMCglmm on tarsus length
    # fixed effects are sex and inbreeding
    # random effects are the animal (VA)
    # and the ID (VPE)
    
    model <- MCMCglmm(tarsus ~ 1+sex+f, random = ~animal+ID,
                      pedigree = gen, data = pheno, prior = prior,
                      nitt = 11000, thin = 10, burnin = 1000, verbose = T)
    
    # extract results from MCMCglmm
    posterior.heritability <- model$VCV[,"animal"]/
      (model$VCV[,"animal"]+model$VCV[,"units"]+model$VCV[,"ID"])
    h[k, j] <- mean(posterior.heritability) 
    se_h[k, j] <- sd(posterior.heritability)
    VA[k,j] <- mean(model$VCV[,"animal"])
    VE[k,j] <- mean(model$VCV[,"units"])
    VPE[k,j] <- mean(model$VCV[,"ID"])
    
  }
  
}


  # compute means across simulations
  h_ave <- rowMeans(h)
  VA_ave <- rowMeans(VA)
  VE_ave <- rowMeans(VE)
  VPE_ave <- rowMeans(VPE)
  se_h <- rowMeans(se_h)

  # merge all values with naive ones
  lambda <- c(0.17, lambda0)
  h_ave <- c(h0, h_ave)
  VA_ave <- c(VA0, VA_ave)
  VE_ave <- c(VE0, VE_ave)
  VE_ave <- c(VPE0, VPE_ave)
  se_h <- c(s0, se_h)

  # Extrapolation phase of PSIMEX
  p.names <- c("h",  "se")
  estimates <- data.frame(h_ave,  se_h)
  colnames(estimates) <- p.names
 
    #linear case

    extrapolation_h<- lm(estimates[ ,1] ~ lambda)
    h_pred<-predict(extrapolation_h, newdata = data.frame(lambda = 0))

    extrapolation_se<- lm(estimates[ ,2] ~ lambda)
    se_pred<-predict(extrapolation_se, newdata = data.frame(lambda = 0))

    #quadratic case
  
    extrapolation_h1<- lm(estimates[ ,1] ~ lambda+ I(lambda^2))
    h_pred1<-predict(extrapolation_h1, newdata = data.frame(lambda = 0))

    extrapolation_se1<- lm(estimates[ ,2] ~ lambda+ I(lambda^2))
    se_pred1<-predict(extrapolation_se1, newdata = data.frame(lambda = 0))
    
    #cubic case
    
    extrapolation_h2<- lm(estimates[ ,1] ~ lambda+ I(lambda^2)+ I(lambda^3))
    h_pred2<-predict(extrapolation_h2, newdata = data.frame(lambda = 0))
    
    extrapolation_se2<- lm(estimates[ ,2] ~ lambda+ I(lambda^2)+ I(lambda^3))
    se_pred2<-predict(extrapolation_se2, newdata = data.frame(lambda = 0))
    
    
    # calculate SIMEX SE
    
    # first component 
    # sampling variability
    S <- c()
    #one per lambda 
    for ( i in 1:length(lambda0)) {
        diff <- c()
        #calculate the differences per each simulation
        for ( j in 1: nsim ) {
              diff[j] <- h[i,j]-h_ave[i]
      }
  diff <- na.omit(diff)
  S[i] <- 1/(nsim-1)*sum(diff^2) 
  
}
S <- c(se_h[1], S)

#extrapolated value

#linear case
#
extrapolation_var<- lm(S ~ lambda)
var_pred<-predict(extrapolation_var, newdata = data.frame(lambda = 0))

#quadratic case

extrapolation_var1<- lm(S ~ lambda+ I(lambda^2))
var_pred1<-predict(extrapolation_var1, newdata = data.frame(lambda = 0))

#cubic case

extrapolation_var2<- lm(S ~ lambda+ I(lambda^2)+ I(lambda^3))
var_pred2<-predict(extrapolation_var2, newdata = data.frame(lambda = 0))


# TOTAL SE
#linear 
var_ext <- se_pred^2 + var_pred

#quadratic
var_ext1 <- se_pred1^2 + var_pred1


#cubic
var_ext2 <- se_pred2^2 + var_pred2

true <- h_true
true_low <- true - 1.96*se_true
true_up <- true + 1.96*se_true

naive <- h0
naive_low <- naive - 1.96*s0
naive_up <- naive + 1.96*s0

h_pred_low1 <- h_pred - 1.96*var_ext
h_pred_up1 <- h_pred + 1.96*var_ext

h_pred_low2 <- h_pred1 - 1.96*var_ext1
h_pred_up2 <- h_pred1 + 1.96*var_ext1

h_pred_low3 <- h_pred2 - 1.96*var_ext2
h_pred_up3 <- h_pred2 + 1.96*var_ext2

AIC(extrapolation_h)

AIC(extrapolation_h2)

AIC(extrapolation_h1)


  
```
