## Appendix 1 for "The simulation extrapolation technique meets ecology and evolution: A general and intuitive method to account for measurement error"

#### 1 Variance estimator for SIMEX

Variance estimation for the error-free estimate is an essential part of the SIMEX method, but its derivation is not straightforward. Stefanski and Cook (1995) suggested the jackknife estimator in the context of the SIMEX methodology and defined an analogy between these two concepts. The "error-free" estimate provided by SIMEX, denoted as $\hat{\Theta}_{\text{SIMEX}}$, has essentially two components of variability that need to be taken into account and included in the estimator of its variance. Recall that we denote with $k = 1, ..., K$ the iteration number in the simulation phase, with $\zeta$ the error proportion and with $\hat{\Theta}_k(\zeta)$ the value of the estimated parameter at iteration $k$ for error proportion $\zeta$.

The **first component of variance** is the variance of the error-free estimate itself, due to sampling variability. It is calculated at each simulation step simply as the variance of the estimate given by simulation, $\mathsf{Var}(\hat{\Theta}_k(\zeta))$. In our specific cases, this is the squared standard error of the paremeter estimates of the regression model used to calculate inbreeding depression or the animal model used to compute heritability. This quantity is then averaged over the total number of simulations $K$ for a fixed error proportion $\zeta$, given as $\mathsf{Var}(\hat{\Theta}(\zeta)) = K^{-1} \sum_{k=1}^{K} \mathsf{Var}(\hat{\Theta}_k(\zeta))$. An extrapolation is then performed for these variances as well, using the same function selected for the extrapolation of the mean, which provides an estimate of the sampling variance corresponding to an error proportion of zero, $\mathsf{Var}(\hat{\Theta}(\zeta = 0))$.

The **second component of variance** is due to the measurement error variability, *i. e.* the error $\Delta_k(\zeta) = \hat{\Theta}_k(\zeta) - \hat{\Theta}(\zeta)$ that occurs in averaging across $K$ simulations, and is what establishes the connection with the jackknife methodology. An approximate value of this component can be calculated for each value of the error proportion $\zeta$ from the differences between the estimate given in each simulation step and the global estimate obtained by averaging:

$$s_{\Delta}^{2}(\zeta) = \frac{1}{K-1} \sum_{k=1}^{K} (\hat{\Theta}_{k}(\zeta) - \hat{\Theta}(\zeta))^{2} \quad (1)$$

where $\hat{\Theta}_k(\zeta)$ is the SIMEX estimate of iteration $k$ for error proportion $\zeta$, and $\hat{\Theta}(\zeta)$ is the mean of these $K$ iterations. After calculating (1) for each error $\zeta$, the extrapolation phase is repeated to obtain an estimate for the error-free value $s_{\Delta}^{2}(\zeta = 0)$.

The **total variance** is the sum of these two components, $\mathsf{Var}(\hat{\Theta}_{\mathrm{SIMEX}}) = \mathsf{Var}(\hat{\Theta}(\zeta = 0)) + s_{\Delta}^{2}(\zeta = 0)$, and it is reported together with the error-free estimate provided by SIMEX.

### 2 Misassigned paternity effects in song sparrows

#### 2.1 Inbreeding

The percentage of misassigned paternities among birds for which we have records of both tarsus length and juvenile survival is 17%, calculated as the discrepancy between apparent and actual fathers. The difference among inbreeding coefficients of each bird calculated from the two different pedigrees is denoted as $d_i = f_i^{(\text{apparent})} - f_i^{(\text{actual})}$ and its distribution is shown in Figure S1. The distribution is clearly non-Gaussian, with overproportional weight on 0 and comparably long tails. Moreover, Figure S2 illustrates that the error $d_i$ depends on the actual inbreeding coefficient $f_i^{(actual)}$, due to the fact that the inbreeding coefficient cannot be negative.

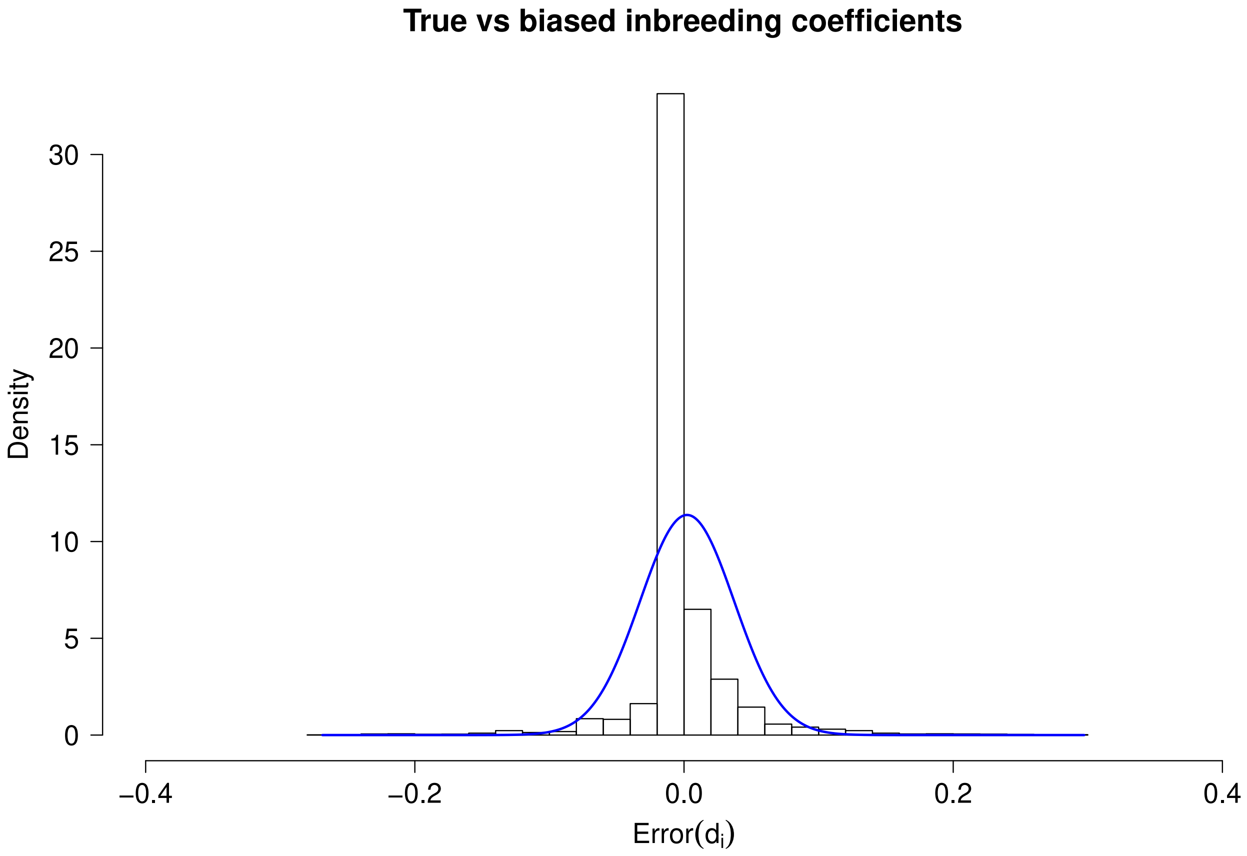


Figure S1: Error distribution of inbreeding coefficients as deduced from the apparent song sparrow pedigree. The blue line represents a normal distribution with the same mean and standard deviation as the $d_i$ values, underlining the non-normality of the error.

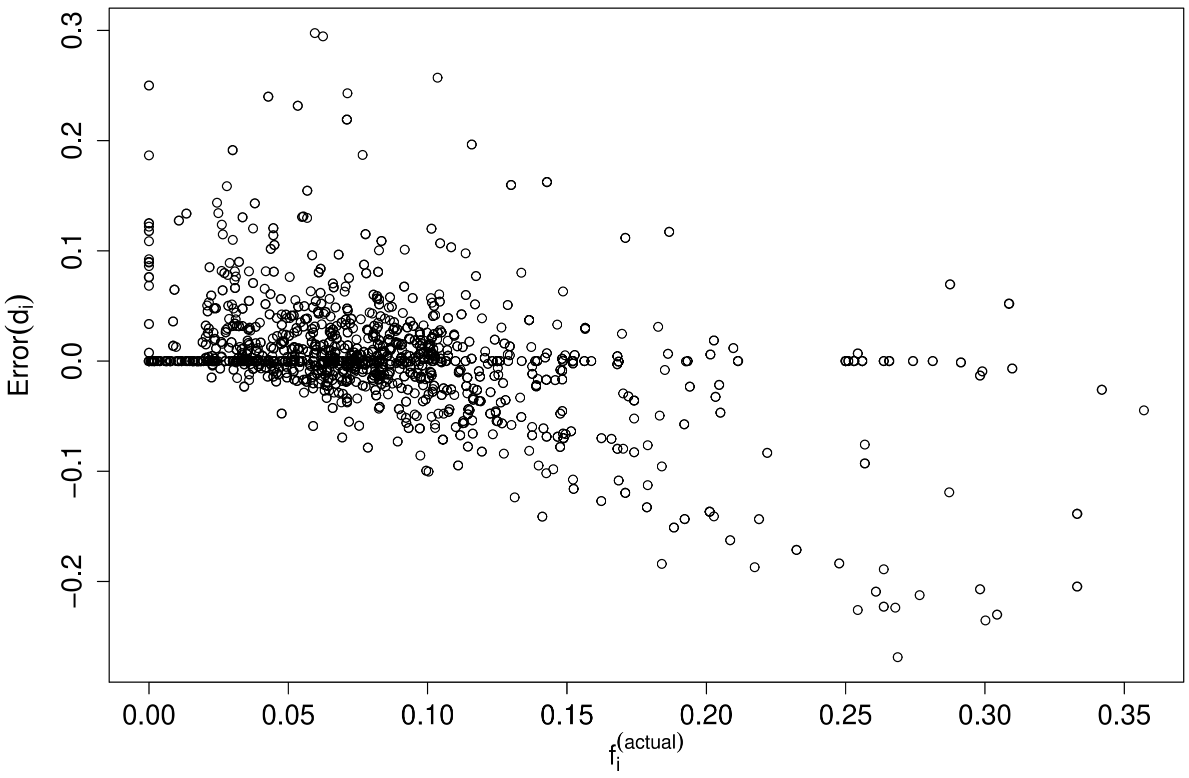


Figure S2: Error of the inbreeding coefficients in the apparent song sparrow pedigree ($d_i$) as a function of the correct inbreeding coefficients from the actual pedigree ($f_i^{(\mathrm{actual})}$).

#### 2.2 Variance components of heritability

The model used in the main text to estimate the heritability of tarsus length in song sparrows was

$$y_{ij} = \mu + \beta_f f_i + \beta_{\text{sex}} sex_i + a_i + id_i + R_{ij}$$

The model decomposes variance into additive genetic variance $\sigma^2_A$, permanent environmental variance $\sigma^2_{PE}$ and residual variance $\sigma^2_R$, and heritability is calculated as $h^2 = \sigma^2_A/(\sigma^2_A + \sigma^2_R + \sigma^2_{PE})$. The posterior distributions of these variances from the MCMC procedure are shown in Figure S3, both for the case when the calculations were based on the genetic (actual) and the error-prone (apparent) pedigree.

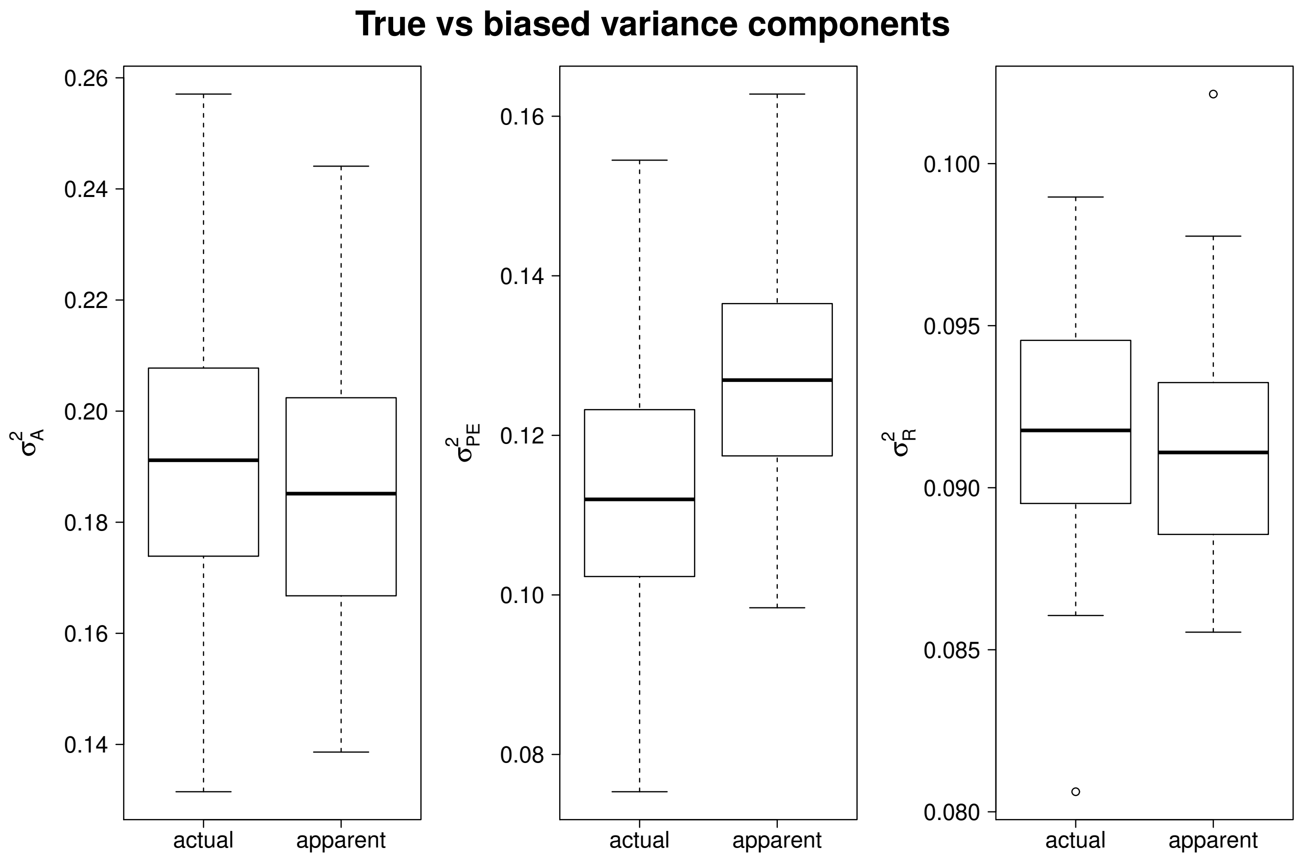


Figure S3: Posterior distributions of the variance components from the MCMC samples used to calculate heritability of tarsus length in song sparrows from the actual and apparent pedigrees.

### 3 R Code for simulations

The `R` code for the simulation of pedigrees is given in the `R` scripts in `Appendix2.R` and `Appendix3.R`. Examples of the application of the PSIMEX algorithm on error prone data are reported in `Appendix4.R` and `Appendix5.R`, while the code for the application to the song sparrow dataset is given in `Appendix6.R` and `Appendix7.R`. To implement PSIMEX on already existing error prone pedigrees, please also refer to the vignette of the `R` package `PSIMEX` (Ponzi, 2017).

### 4 Starting with the wrong error proportion

As discussed in the main text, accurate knowledge of the true error proportion in the pedigree is required to apply error correction via PSIMEX. In this section we illustrate how the error-corrected estimate $\hat{\Theta}_{\text{SIMEX}}$ is sensitive to appropriate knowledge of the true error proportion, denoted here as initial error proportion $\zeta_I$, when calculating error-corrected estimates for inbreeding depression. To this end, we start with a range of wrong initial misassigned paternity error proportions on a simulated pedigree. In addition, we use two wrong initial values for $\zeta_I$ in the example of the song sparrow pedigree, which were larger and lower than the actual initial error proportion.

#### 4.1 Simulations

We considered a single simulated pedigree with $Ne/Nc = 1$ and an initial error proportion of 0.17. PSIMEX with $K = 100$ iterations was then applied to estimate inbreeding depression assuming initial error proportions between 0 and 0.5 and starting from a $\beta_f$ in the error-free pedigree equal to -6.5. Figure S4 illustrates how PSIMEX tends to recover an over-corrected (thus overestimated) “error-free” $\hat{\Theta}_{\text{SIMEX}}$ value when assuming an initial error proportion that is larger than the actual error. This is intuitively reasonable, because it is then implicitly assumed that the error-prone estimate is a result of more error and therefore a larger correction is expected. Since the expected trend for a pedigree with $Ne/Nc = 1$ is an increasing value of inbreeding depression, it is expected that an over-correction will result in lower estimates of inbreeding depression. On the other hand, when starting from an initial error proportion that is too small, PSIMEX tends to under-correct the error-prone estimate of inbreeding depression and retrieve a higher value of inbreeding depression.

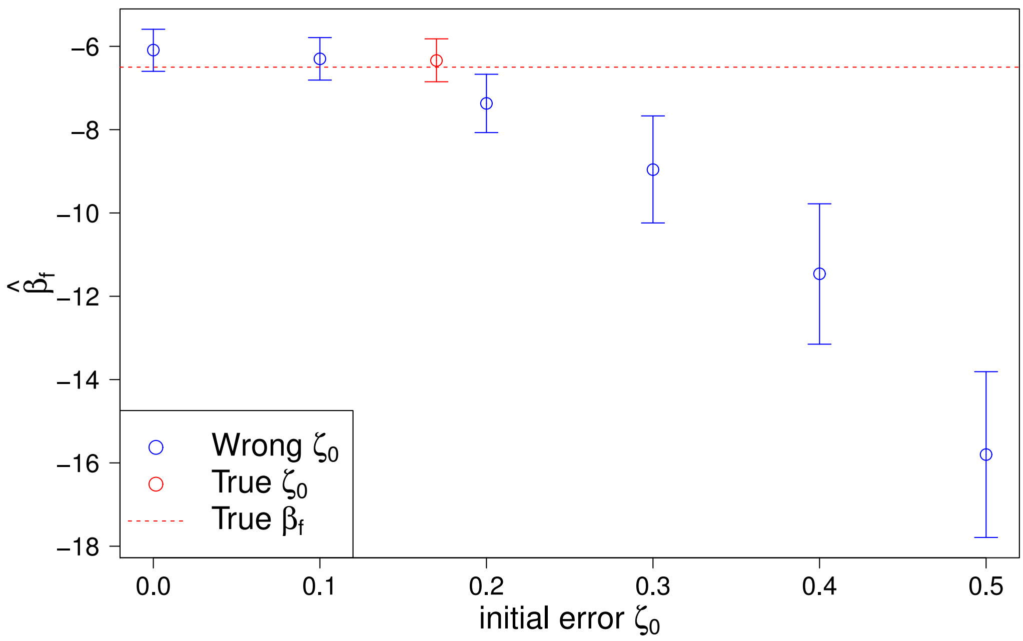


Figure S4: PSIMEX estimates of inbreeding depression in a simulated pedigree, depending on the initial error proportion $\zeta_I$ used for the procedure. Red indicates the estimate when assuming the correct initial proportion, and the dotted line marks the actual value. The extrapolation function with the lowest AIC was used in all cases and resulted in the quadratic function in most cases. The estimates are reported together with their 95% confidence intervals.

#### 4.2 Song sparrows

Using the same setup and number of iterations $K = 100$ as in the main text, it was first assumed that the initial error proportion was $\zeta_I = 0.35$ instead of the correct $\zeta_I = 0.17$, thus larger than the real error in the song sparrows example. PSIMEX then results in an over-correction of the estimate of inbreeding depression, see Figure S5. On the other hand, when assuming an initial error proportion of $\zeta_I = 0.08$, the correction is too small (Figure S6).

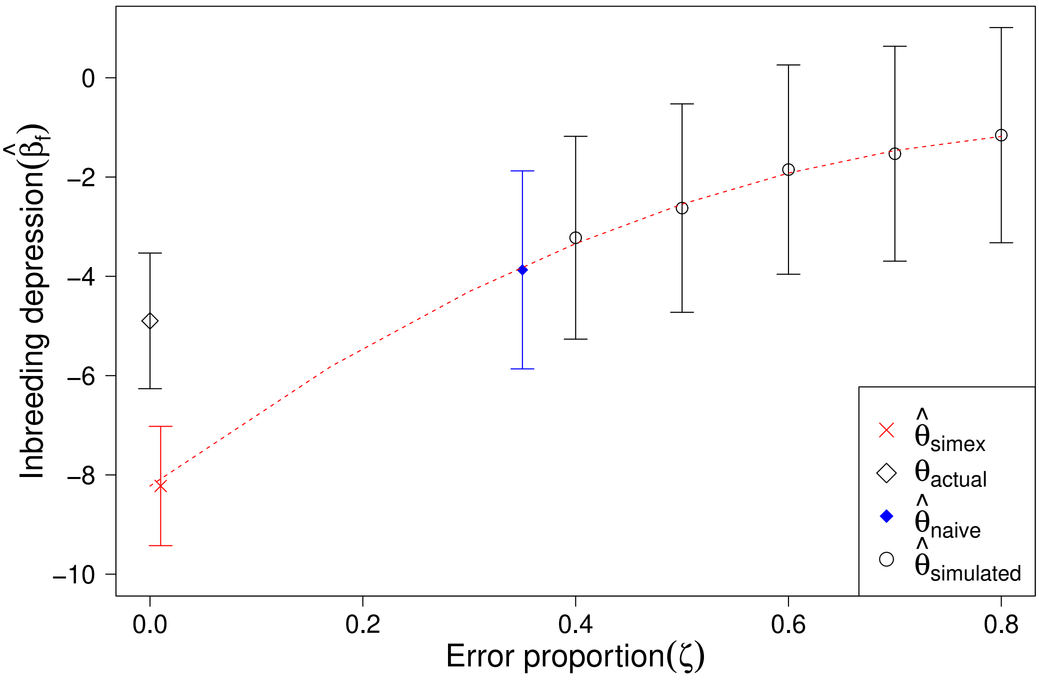


Figure S5: Simulation and extrapolation phase of the PSIMEX procedure to recover estimates of inbreeding depression in the song sparrows when the initial error proportion was erroneously set to $\zeta_I = 0.35$.

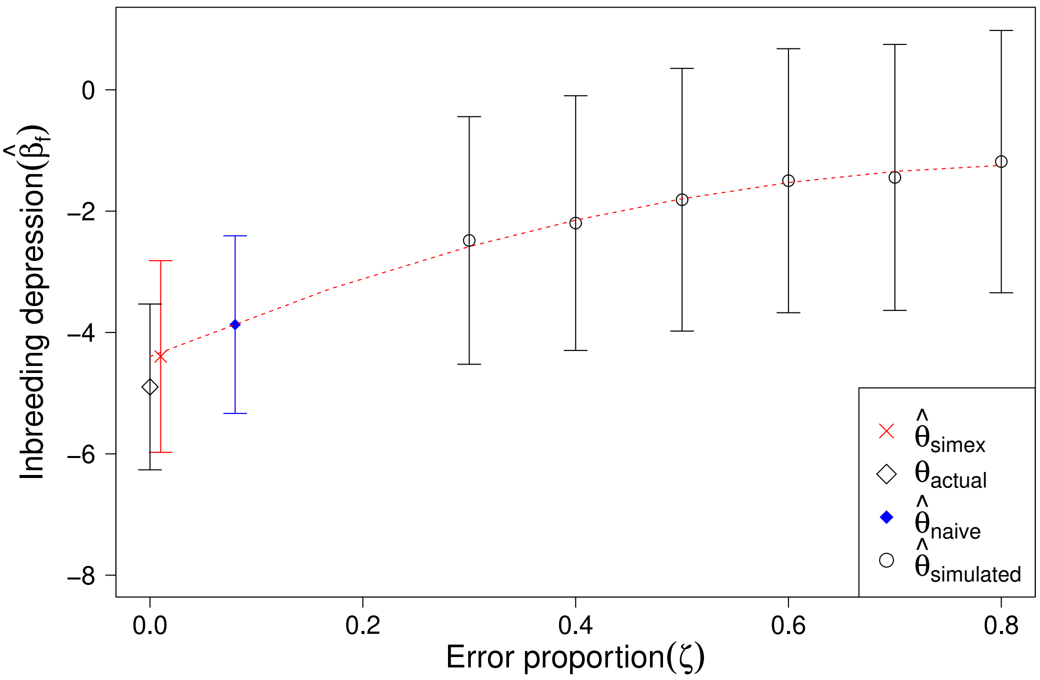


Figure S6: Simulation and extrapolation phase of the PSIMEX procedure to recover estimates of inbreeding depression in the song sparrows when the initial error proportion was erroneously set to $\zeta_I = 0.08$.

### 5 Starting with the wrong error-generating mechanism

It is not only important to start from an accurate error proportion, but also to use an accurate error-generating mechanism in the simulation phase of PSIMEX. This section illustrates how estimates can be biased if assumptions about the error-generating mechanism in the pedigree are wrong. Using the song sparrows dataset as an example, we first ran PSIMEX erroneously assuming that misassigned paternities only occurred in the first (older) half of the pedigree (Figure S7) and second only in the second (younger) half of pedigree (Figure S8). Third, we erroneously assumed that females mate with similar males and extra pair paternities are therefore not at random, but with individuals that are similar to the real father. Thus PSIMEX was implemented assuming that misassigned fathers are similar to the genetic father in terms of a specific trait (here body mass). Instead of replacing a father with a random individual from the same generation, we replaced him with the individual in the same generation who had the most similar body mass (Figure S9).

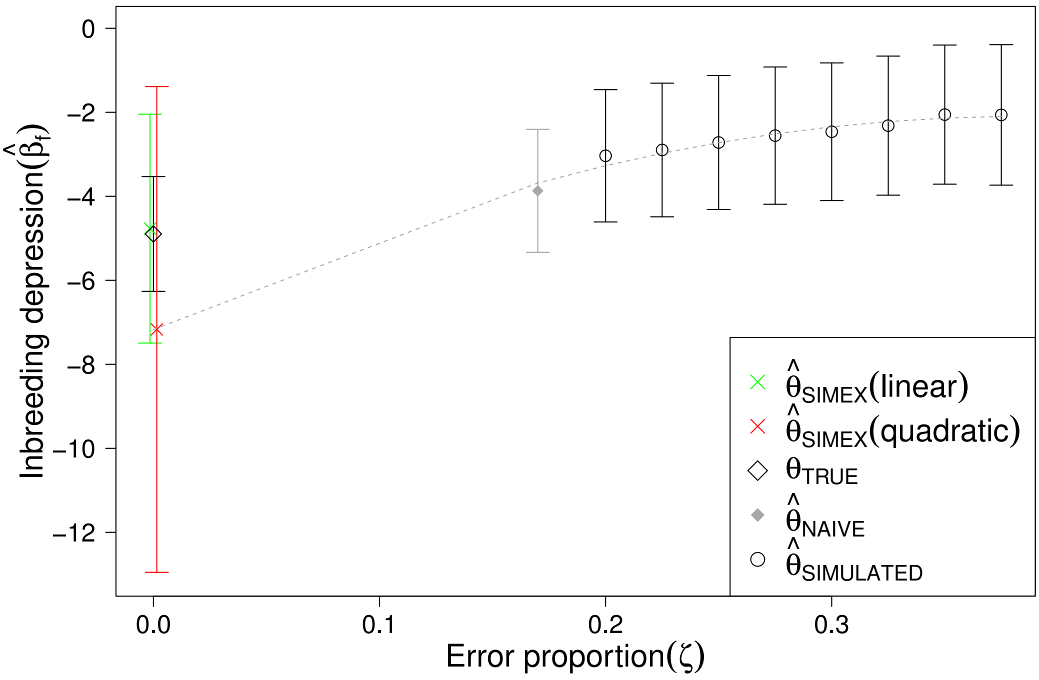


Figure S7: PSIMEX results for inbreeding depression in song sparrows when the error was erroneously assumed to occur only in the first half of the pedigree. The $x$-axis represents the overall error proportion in the complete pedigree, thus the upper limit is 0.5 given that errors only occurred in the first half of the pedigree.

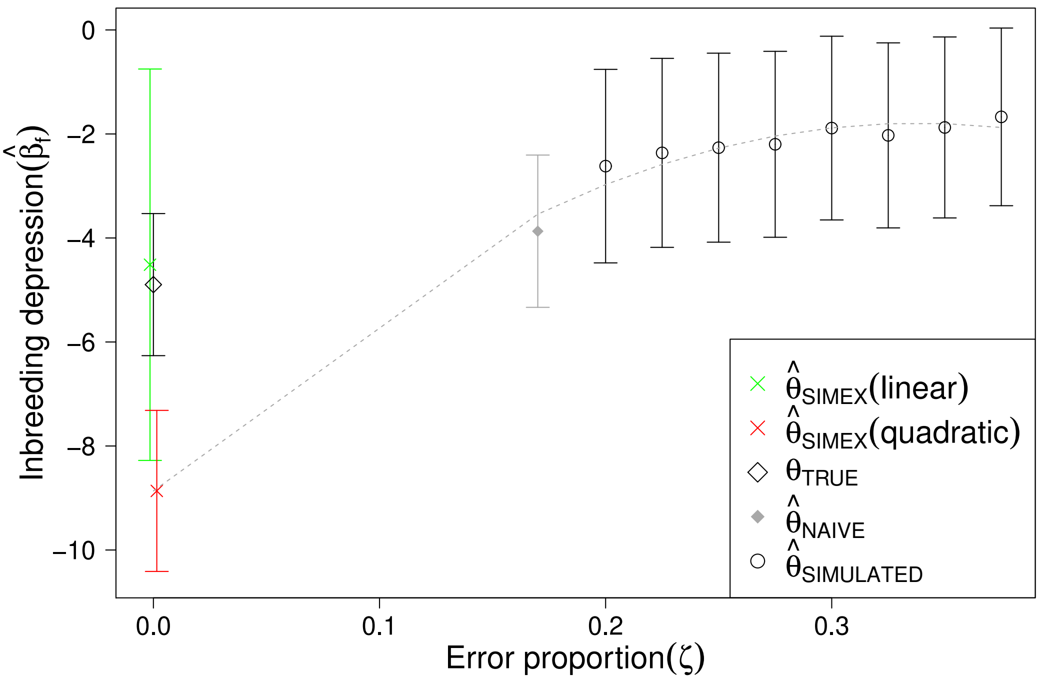


Figure S8: PSIMEX results for inbreeding depression in song sparrows when the error was erroneously assumed to occur only in the second half of the pedigree. As in Figure S7 the $x$-axis represents the overall error proportion in the complete pedigree, thus the upper limit is 0.5.

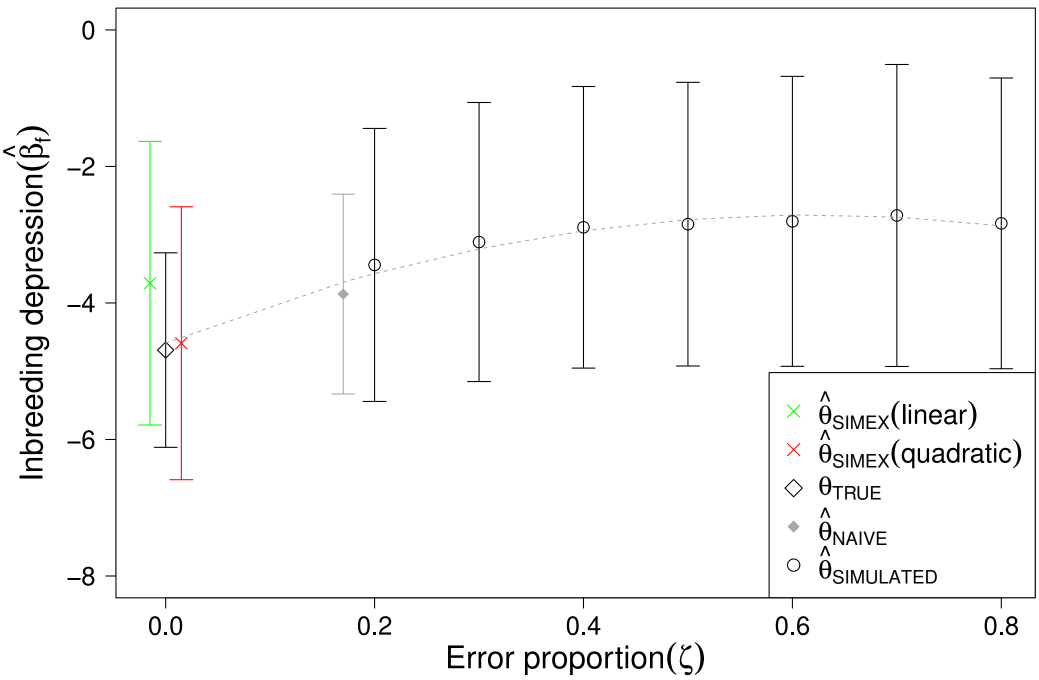


Figure S9: PSIMEX results for inbreeding depression in song sparrows when the replacement of fathers is assumed to be with phenotypically similar individuals.

These examples illustrate how assuming the wrong error generating mechanisms can lead to biased PSIMEX estimates, with extrapolated estimates lower or higher than the true values. Thus, assumptions about the error-generating mechanism need to be validated with data to be able to properly implement the simulation and the extrapolation phases of PSIMEX. It also points out the importance of the choice of the extrapolation function, with different functions leading to different estimates of the error-corrected parameters.

### References

Ponzi, E. (2017). *PSIMEX: SIMEX Algorithm on Pedigree Structures.* University of Zurich. R package version 1.0.

Stefanski, L. and J. Cook (1995). Simulation-extrapolation: The measurement error jackknife. *Journal of the American Statistical Association 90*, 1247–1256.
